# Kinetic properties of optogenetic site-specific DNA recombination by LiCre-*loxP*

**DOI:** 10.1101/2024.05.17.594525

**Authors:** Alice Dufour, Hélène Duplus-Bottin, Thomas Boukéké-Lesplulier, Eliane Casassa, Gérard Triqueneaux, Léo Tarbouriech, Camille Darthenay-Kiennemann, Agnès Dumont, Catherine Moali, Franck Vittoz, Daniel Jost, Gaël Yvert

## Abstract

Advances in optogenetics now allow to specifically modify the DNA of live cells with light. However, using these technologies successfully requires to know their properties in terms of sensitivity, efficiency, kinetics and mechanism. We previously developed an optogenetic tool made of a single chimeric protein called LiCre that enables the induction of specific changes in the genome with blue light via DNA recombination between *loxP* sites [1]. Here, we used *in vitro* and *in vivo* experiments combined with kinetic modeling to provide a deeper characterization of the photo-activated LiCre-*loxP* recombination reaction. We find that LiCre binds DNA with high affinity in absence of light stimulus and that this binding is cooperative although not as much as for the Cre recombinase from which LiCre was derived. In yeast, addition of riboflavin to the culture medium had no effect on LiCre’s efficiency, even when cells over-expressed riboflavin kinase, suggesting that abundance of the flavin mono-nucleotide co-factor is not limiting for the reaction. However, LiCre’s efficiency in yeast gradually increased when raising temperature from 20°C to 37°C. The recombination kinetics observed in live cells are best explained by a model where photo-activation of two or more DNA-bound LiCre units (happening in seconds) can produce (in several minutes) a functional recombination synapse. This model was able to capture the effect of a point mutation altering LiCre’s light cycle. This deeper understanding of the LiCre-*loxP* system provides additional knowledge for designing experiments where specific genetic changes are induced in live cells with light.

**AUTHOR SUMMARY:** Using a technology called optogenetics, scientists are now able to change the DNA sequence of live cells by illuminating them with light. In theory, they can trigger in genetically-engineered organisms a mutation of interest in specific cells at a specific time. This practice, however, is not common because optogenetics relies on light-controlled enzymes that are recent and not well characterized. We previously developed one of these enzymes, called *LiCre*, which recognizes a specific piece of DNA. Following illumination with blue light, *LiCre* can switch on or off whatever gene is close by. Here, we combined experiments and computational modeling to better understand how, and how fast, *LiCre* works. We find that, although it is inactive in the dark, *LiCre* does not need photo-activation to bind to its DNA partner. We estimated the speed at which *LiCre* gets deactivated in the dark and the speed at which active *LiCre* molecules modify DNA. We also showed the effect of temperature and illumination dynamics on *LiCre*’s efficiency. These results will help design strategies where *LiCre* can be used to conduct genetic studies at high spatio-temporal resolution, or implement it in industrial and biomedical applications.

## INTRODUCTION

Experimental biologists usually make discoveries by perturbing living organisms and observing the consequences of these perturbations. Precision in these manipulations is often crucial because biological systems are spatially complex and highly dynamic. Optogenetics offers the possibility to alter cellular activities with light and provides unprecedented spatio-temporal resolution. For this reason, a wide set of light-inducible biotechnologies have been developed in the past years. These tools include photo-activatable ion channels that trigger neuronal activities when illuminated (see [2] for a review on neuroscience applications), photo-controlled kinases activating signaling pathways, as well as systems controlling protein localization, protein-protein interactions, gene expression or DNA editing (see [3] for a review on molecular tools). Many of these systems are genetically encoded, thereby alleviating the need for chemicals. They are all based on conformational changes of a macromolecule leading to its activation or inactivation in response to light. Yet, each tool has its own specificity in terms of light sensitivity, efficiency and response dynamics; optimal usage of a given tool therefore requires quantitative knowledge of its biochemical properties.

We recently developed an optogenetic site-specific DNA recombinase called LiCre [1]. As for the Cre recombinase from which it derives, LiCre recognizes 34-bp DNA sequences called *loxP* and catalyzes DNA recombination between two such sites. This allows experimentalists to induce specific changes in the genome of live cells in real time with blue light. For example, if a gene is flanked by two *loxP* sites in LiCre-expressing cells, shining light on these cells can delete the gene. Compared to other photo-activatable recombinases which are based on two protein halves (splitCre) that complement in response to light [4–7], LiCre is unique as it consists in a single chimeric protein comprising the Light-Oxygen Voltage domain from *Avena sativa* (asLOV2) fused to a mutant version of Cre. It was designed such that the unfolding of a critical α-helix in response to light enables the assembly of a functional recombination synapse [1]. The mode of action of LiCre, represented in Fig. 1a, is speculated from the abundant knowledge previously acquired on the Cre-*loxP* synapse [8] and the asLOV2 photo-switch [9]. Formation of the Cre-*loxP* complex involves the cooperative binding of two Cre units on each *loxP* site and the subsequent assembly of the four Cre proteins complexed with their DNA substrates [8]. The resulting intasome is stabilized by interactions between adjacent protomers: on one side of the DNA substrate, the four C-terminal domains are locked together in a cyclic manner, each unit hosting the αN helix of its neighbor in a nest; on the other side of DNA (depicted in Fig. 1a for LiCre), N-terminal domains also form a cycle of binding interfaces but in the reverse orientation [10]. This way, each protomer is stabilized by its two neighbors. For LiCre, the strength of interactions between C-terminal domains was reduced by mutating two critical residues of the αN helix; the association of N-terminal domains was perturbed by truncating the αA helix interacting with the neighboring unit; and photo-activation was obtained after designing a chimeric α-helix comprising Jα from asLOV2 and the remaining part of αA from Cre [1].

**Figure 1.**
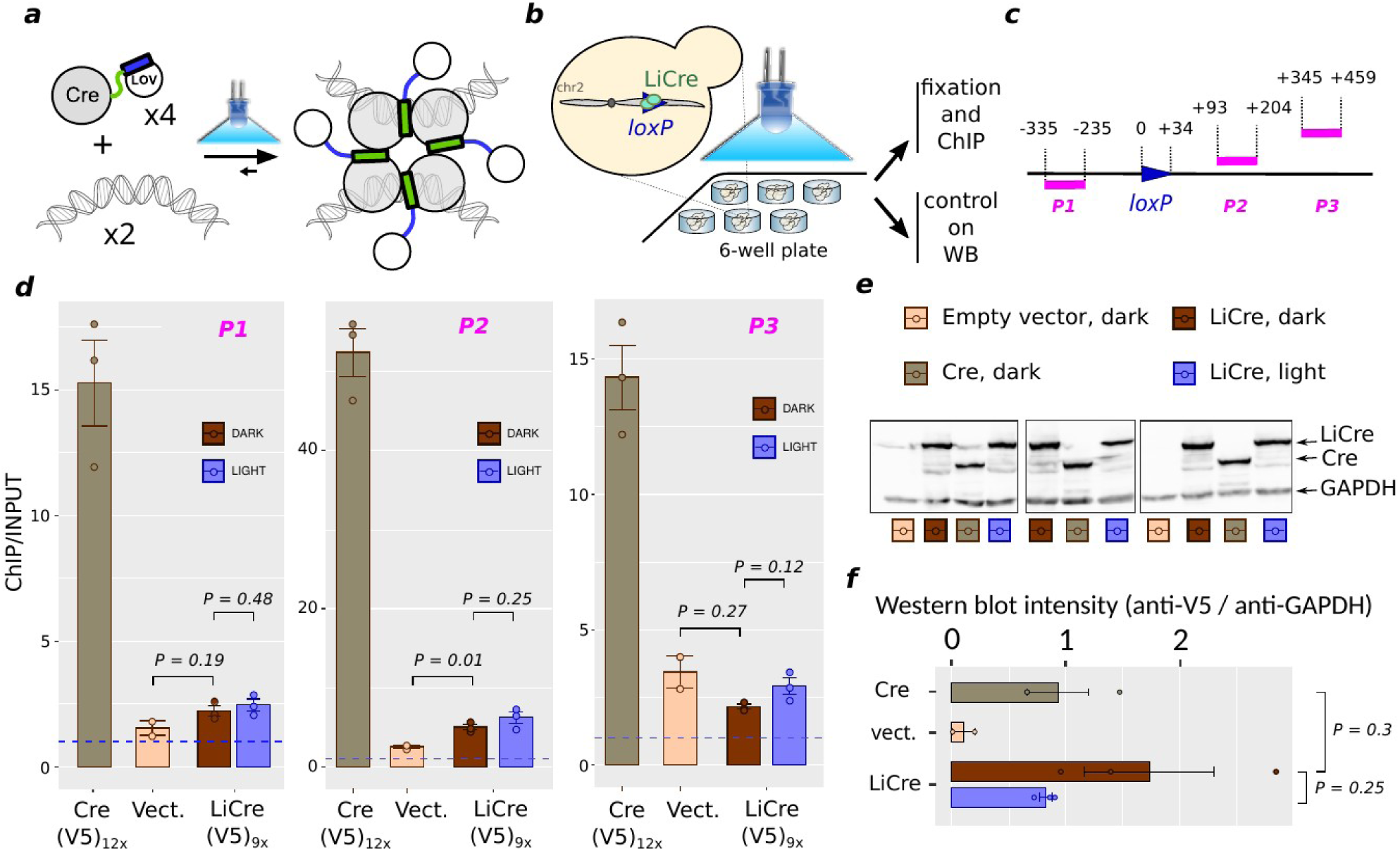
Analysis of LiCre:DNA binding *in vivo*. **a**) Hypothetical principle of LiCre photo-activation. The chimeric α-helix is presented as a rectangle; in blue when folded as the Jα helix of LOV2, in green when folded as the αA helix of Cre. Modified from [1]. **b**) Chromatin immunoprecipitation (ChIP) experimental setup. **c**) Position of qPCR-amplified regions *P1*, *P2* and *P3* relative to *loxP*. **d**) ChIP-qPCR quantifications. Every dot corresponds to one biological replicate (independent culture inoculated from an independent transformant of the Cre-V5 or LiCre-V5 expression plasmid). Values correspond to enrichment by ChIP (relative to its input), normalized by the signals obtained on three unrelated loci of the genome (see Methods). *P* values: Welch *t*-test. **e**) Western blot analysis of whole-cell protein extracts. Proteins were extracted from the same population of cells as for ChIP. **f**) Quantification of LiCre, Cre and GAPDH immunoblot shown in e. *P* values: Welch *t*-test.

Although LiCre outperformed split-based systems in terms of speed and efficiency [1], two essential questions remain on its properties. First, it is unclear if its association to *loxP* occurs before or after its photo-activation. Second, the observed dependency of its efficiency on the dynamics of the light stimulus is not explained. Here, we address these questions by combining *in vitro* and *in vivo* experiments with kinetic modeling. We found that LiCre units bind to DNA with high affinity even if they are not photo-activated; that they do so in a cooperative manner although this cooperativity is weaker than for Cre; that temperature greatly affects LiCre efficiency, that the kinetics of recombination in live cells are best explained by a model where photo-activation of two or more DNA-bound LiCre units happens in seconds and enables the formation, in several minutes, of a functional recombination synapse; and we identify a point mutation in LiCre that modifies its light cycle.

## RESULTS

### LiCre binds its target DNA in absence of photo-stimulation

LiCre photo-activation is based on a conformational change that enables protein-protein interactions assembling the recombinogenic intasome [1]. We previously proposed two possible models for this activation. In the first model, LiCre monomers are unable to bind DNA *loxP* sites in their ‘dark’ – inactive - state and the photo-induced conformational change restores this ability. In the alternative model, LiCre binds to *loxP* sites even in the dark, and the conformational change of photo-stimulated DNA-bound LiCre molecules enables the assembly of the recombination synapse. To distinguish between these two models, we evaluated the ability of LiCre to bind DNA in conditions where it was not photo-activated.

We first tested this DNA-binding ability *in vivo* by using chromatin immuno-precipitation (ChIP). We constructed yeast strains harboring a single *loxP* site in the genome and expressing LiCre or Cre tagged with repeated V5 epitopes (Fig. 1b). We cultured cells overnight and then either let them in dark conditions or, for LiCre-V5 samples, illuminated them in conditions known to trigger LiCre activity [1]. Following illumination, we processed half of the cells for fixation and ChIP using an anti-V5 antibody, and the other half for Western blot. We quantified ChIP by qPCR at three positions near the *loxP* site (Fig. 1c) and, for normalization, at three unrelated positions on the genome. As shown on Fig. 1d, we observed a strong ChIP signal for all three probes of the *loxP* locus for Cre, consistent with the known high-affinity of Cre for its *loxP* target [8]. Regarding LiCre, we observed a much weaker ChIP signal, which reached statistical significance only for the probe located very close to *loxP* (probe P2 on Fig. 1c-d). There are several possible explanations for the difference in ChIP signals between LiCre and Cre. First, given that ChIP is a crude assay where fragmented particles of cross-linked material are pulled down using antibodies, its output signal could vary if the two proteins differ in their sub-cellular distribution (eg. nuclear/cytoplasmic ratio), in the way cross-linking affects the accessibility and/or affinity of the antibody (conformation, local molecular crowding), or in their degree of non-specific interactions with cellular factors. Second, expression levels of the two proteins may differ. Third, when verifying the number of V5 epitopes by Sanger sequencing, we found that they had unfortunately amplified for Cre-V5 (12 copies) but not for LiCre-V5 (9 copies, as expected). It is possible that this increased number of epitopes in Cre-V5 contributed to its higher ChIP signal. However, Western-Blots showed that, in denaturing conditions, the detection of Cre-V5 by the anti-V5 antibody was not stronger than the detection of LiCre-V5 (Fig. 1e-f). This rules out a higher expression level for Cre-V5 than LiCre-V5 but does not exclude the possibility of different immuno-affinities in pulled-down conditions. Comparing the DNA affinities of LiCre and Cre therefore requires complementary experiments (provided below). Very importantly, illumination prior to chromatin immuno-precipitation had clearly no effect on LiCre ChIP signals (Fig. 1d). We conclude that, *in vivo*, LiCre binds *loxP*-containing chromatin in the dark and that this binding is equally strong in light conditions.

To better quantify LiCre:DNA binding, we analyzed this affinity *in vitro* by surface plasmon resonance (SPR). This method offers high sensitivity to monitor macromolecular associations and dissociations; it was previously used to characterize the binding affinity of Cre to *loxP* [11]. We produced and purified recombinant Cre and LiCre proteins from *E. coli* (Fig. 2a-b) and injected these proteins into microfluidic chips containing immobilized biotinylated DNA substrates (see Methods). We used DNA substrates containing either a full *loxP* site that can associate with two units of Cre or LiCre, or a half site allowing the binding of only one unit. We also used a randomized DNA sequence to quantify non-specific binding. We did not apply any blue light illumination to the samples, neither before injection of the protein solution in the chip nor during acquisitions.

**Figure 2.**
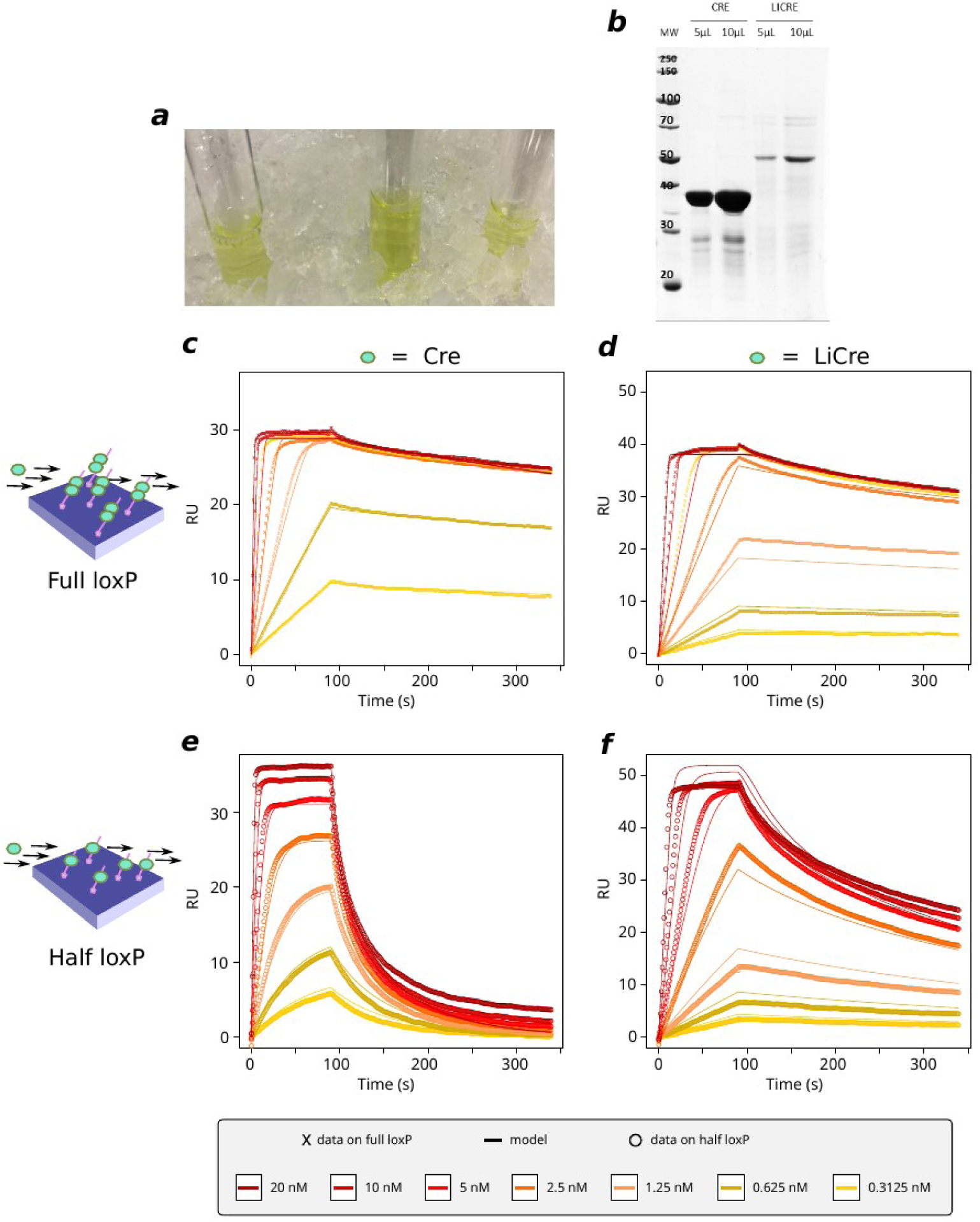
Analysis of LiCre:DNA interaction *in vitro* by surface plasmon resonance (SPR). **a)** Glass tubes containing eluted fractions of recombinant LiCre protein purified from *E. coli.* **b)** SDS-PAGE analysis of purified recombinant Cre and LiCre proteins. **c-f)** Net sensorgrams showing Cre (**c,e**) or LiCre (**d,f**) interaction with full (**c,d**) or half (**e,f**) *loxP* sites. At time zero, recombinant proteins were injected at the indicated concentrations over a BIACORE sensor chip containing immobilized *loxP* sites (c,d), *loxP* half sites (e,f) or unrelated DNA as negative control blank. After 90 seconds, the sensor chip was washed by injecting running buffer lacking the protein (dissociation step). Net sensorgrams were generated from the raw data by subtracting the negative blank control to correct injection-related fluctuations. Dots, observed values. Lines, predicted values from models (see Text and Methods). Model parameters for Cre (panels c and d) were: *α* = 0.87, *RUmax_h_* = 38.2, *RUmax_f_* = 28.8, *k_t_* = 0.51, *k_1_* = 0.962, *k_-1_* = 1.106, *k_2_* = 0.498, *k_-2_* = 0.000452, which gave a fit score of 0.0011. Model parameters for LiCre (panels d and e) were: *α* = 1.71, *RUmax_h_* = 53.2, *RUmax_f_* = 38.1, *k_t_* = 0.099, *k_1_* = 0.98, *k_-1_* = 0.481, *k_2_* = 4.655, *k_-2_* = 0.00286, which gave a fit score of 0.00193.

As shown in Fig. 2c, Cre association with the full *loxP* site was very rapid and its dissociation was extremely slow. This observation is in perfect agreement with previous experiments [11] based on the same DNA substrates as here. For LiCre, association with the full loxP site was slightly slower than for Cre but still very fast while the dissociation was slightly faster (Fig. 2d). This demonstrates that LiCre is able to bind its DNA target site even if it is not photo-activated, which may explain why the blue-light treatment did not increase the detection of LiCre chromatin-binding *in vivo* (Fig. 1).

The profiles obtained with half *loxP* sites are also very informative. In accordance with [11], Cre dissociation from the half site is much faster than its dissociation from the full site (Fig. 2e). Similarly, for LiCre, dissociation from the half site is also faster, but the difference is not as pronounced as for Cre (Fig. 2f). This suggests cooperativity between the two Cre or LiCre units when binding to the full *loxP* site, and that cooperativity of LiCre in the dark may be lower than for Cre.

### LiCre binding to DNA is strong and cooperative

To rationalize our in vitro measurements in terms of binding affinities, we developed a kinetic model of the association and dissociation of LiCre and Cre molecules to DNA, accounting for the SPR protocol (see Methods). Briefly, based on [11], the binding of one molecule to a half-site is described as a simple one-step assembly reaction; while the binding of two molecules to a full site is assumed to happen sequentially: binding of the first molecule is assumed to follow the same kinetics as for a half-site while the second binding event may have different kinetics due to cooperativity mediated by the presence of the already-bound molecule (Fig. 3a). In addition to the association (*k_1_*, *k_2_*) and dissociation (*k_-1_*, *k_-2_*) rates of each step, the model comprises the following parameters that enable its direct confrontation to experimental acquisitions: a coefficient of mass transport in the biosensor chip (*k_t_*), a linear coefficient (*α*) linking the Response Unit (RU) of the SPR experiments to the amount of bound proteins, and maximal RU values (*RUmax_h_* and *RUmax_f_*) when all of the half or full *loxP* sites are fully occupied, respectively. We fitted these parameters by least-squared minimization between predicted and experimental sensorgrams (Fig. 2c-f, see Methods). To study the identifiability of model parameters, we ran the procedure 1,000 times starting from different initial parameter values picked randomly within acceptable ranges. Among the resulting 1,000 trials, 541 and 859 led to high-quality fits for Cre and LiCre, respectively (Fig. 3b-c). The corresponding sets of parameters are shown in Fig. 3d-f. Fitted values of *RUmax_h_*and *RUmax_f_* had very low standard deviations and were therefore very well defined, both for Cre and LiCre (Fig. 3f). Parameters *k_t_* and *α* were each well estimated for LiCre (Fig. 3e). For Cre, their individual estimation was less precise but, as anticipated by their definition, they co-varied and their product was robustly estimated (Fig 3d,f). Similarly, association and dissociation rates of the first binding reaction were not individually identifiable, but their ratio (K_D1_ = *k_-1_/ k_1_, aka* dissociation constant) was (Fig. 3d-f). The reaction rates of the second binding reaction could be identified for Cre but not for LiCre, however their ratio (K_D2_ = *k_-2_/ k_2_*) was precisely estimated for both proteins (Fig. 3d-f). For Cre, these dissociation constants (K_D1_ = 1.15 nM and K_D2_ = 8.8 × 10^−4^ nM) highlight a very strong cooperativity (K_D2_ << K_D1_) and are consistent with those previously reported [11,12] (see Discussion). Importantly, the cooperativity for LiCre s also strong (K_D1_ = 0.49 nM and K_D2_ = 6.1 × 10^−3^ nM, cooperative moment K_D1_/K_D2_ = 81) but much weaker than for Cre (K_D1_/K_D2_ = 1330). This is consistent with the way LiCre was derived from Cre, which involved the destabilization of protein:protein binding both by introducing point mutations in the αN helix required to lock C-terminal domains together, and by truncating and hindering the αA helix engaged in the interaction between N-terminal domains [1] (see Discussion).

**Figure 3.**
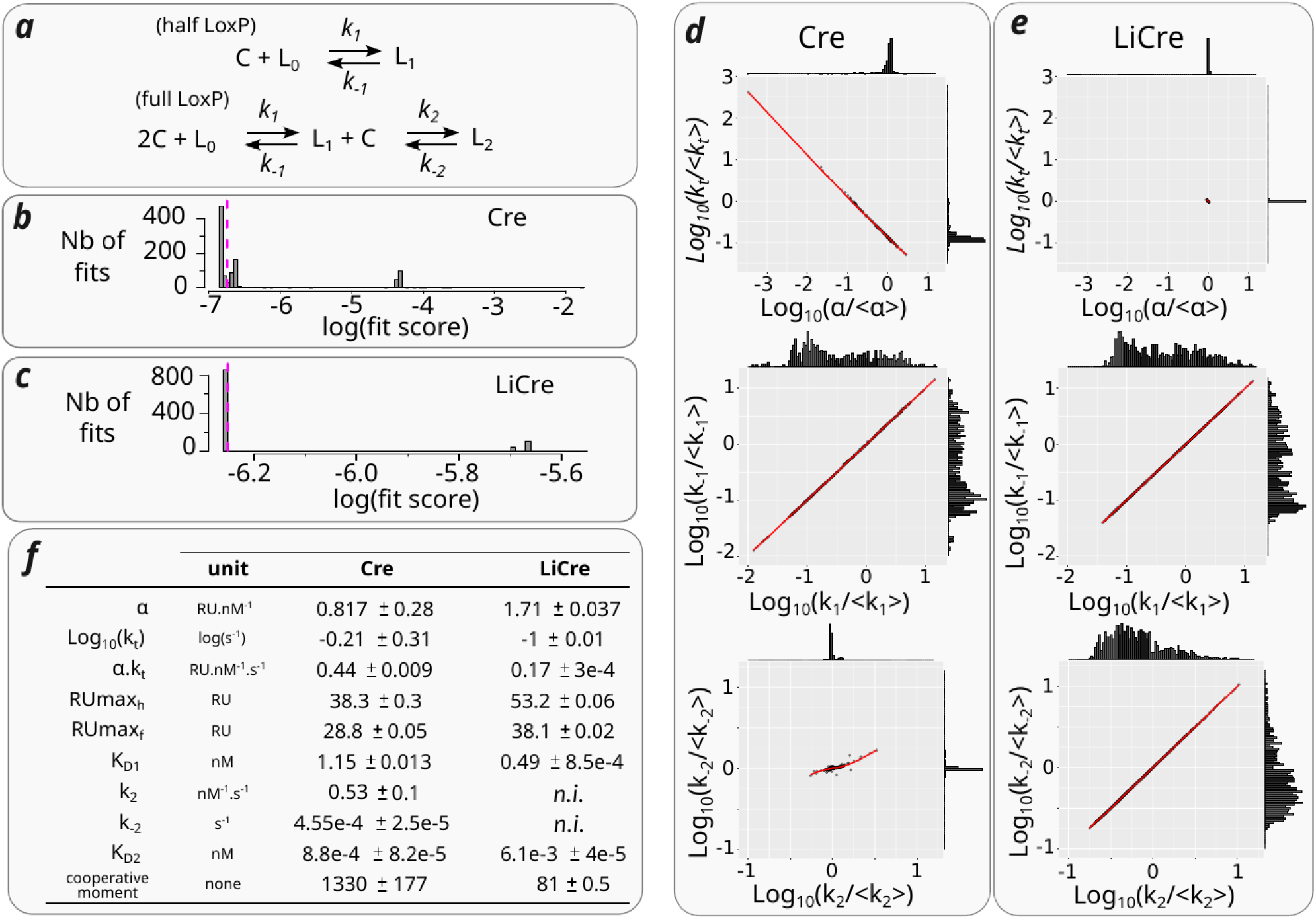
Kinetic modeling of DNA binding. **a)** Model for association of Cre or LiCre molecules (*C*) to DNA *loxP* site (*L*). *L_0_, L_1_* and *L_2_* denote DNA sites occupied by zero, one and two molecules, respectively. **b-c**) Distribution of fit scores across 1,000 optimizations of the DNA-binding model for Cre (b) and LiCre (c). The lower the score, the better the fit (see Methods). Dashed magenta line, arbitrary threshold used to select high-quality fits. **d**) Best estimates of model parameters obtained from 541 optimizations for Cre:DNA binding (see Methods). Every dot corresponds to one optimization. Histograms show marginal distributions of x and y values. Red line, LOWESS local regression. **e**) Same representation as in (d), showing results from 859 optimizations for LiCre:DNA binding. **f**) Table of estimated parameter values (mean +/− sd). *n.i.,* non-identifiable.

### Experimental quantification of LiCre-loxP recombination following various regimes of photo-stimulation

To quantify recombination in live cells, we used a yeast strain harboring a previously-described quantitative reporter [1]. This system is based on the LiCre-mediated excision of a transcriptional terminator preventing the expression of a fluorescent protein (Fig. 4a). This excision occurs by recombination between *loxP* sites located 1.3-Kb from one another. In a typical experiment, a population of yeast cells is illuminated with chosen doses and dynamics of blue light; cells are then re-suspended in fresh medium in the dark to let them express the fluorescent protein if recombination occurred; the population is finally analyzed by flow cytometry to determine the fraction of fluorescent cells. This fraction corresponds to the proportion of stimulated cells that achieved recombination, therefore providing a clear estimate of recombination efficiency. To avoid any potential bias associated with the chromosomal locus where recombination takes place, every cell carried two versions of the reporter, one on chromosome IV coding for mCherry (red fluorescence), and one on chromosome II coding for GFP (green fluorescence). LiCre expression was conferred by a plasmid introduced in the resulting strain. We evaluated the phototoxicity of pulsed blue light on yeast viability by applying illumination to a suspension of cells and by counting colony-forming units (Fig. 4b). This showed that yeast viability is not affected by our illumination conditions.

**Figure 4.**
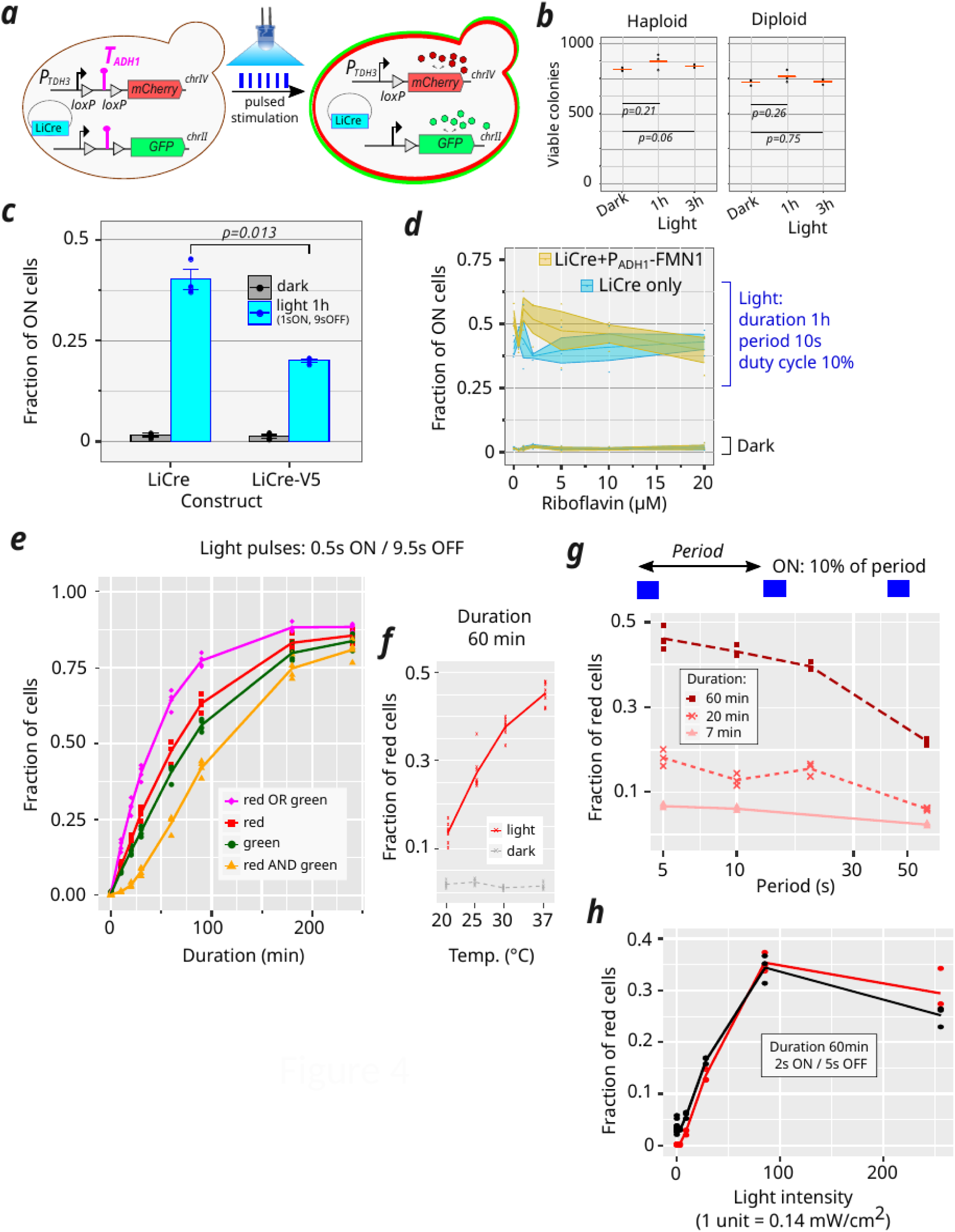
Experimental quantification of *in vivo* LiCre-*loxP* recombination following various photo-stimulation conditions. **a)** Reporter system. Yeast cells harbour two reporter constructs integrated in their genome and express LiCre from a plasmid. A transcriptional terminator (*T_ADH1_*), flanked by *loxP* sites, prevents expression of mCherry or GFP fluorescent proteins. LiCre photo-stimulation triggers recombination, which removes *T_ADH1_* and releases expression. The rate of recombination events can be estimated by counting fluorescent cells in a flow cytometer. **b)** Viability assay of photo-toxicity. Strain GY1761 (haploid) or GY2517 (diploid) was cultivated for 18 hours in three independent cultures (biological replicates), the saturated cultures were transferred to polystyrene flat-bottom well-plates and illuminated or not (dark control) during 1 hr or 3 hrs by pulses of blue light (450 nm, period of 10 s, duty cycle of 10%, intensity of 35 mW/cm2). For each culture, cell dilutions were plated on 3 plates to count colony-forming units (viable colonies). Every dot represents the mean number of colonies on the 3 plates for one biological replicate. Red bar: mean. *P*-value: Welch *t*-test on biological replicates. **c)** Recombination rate induced by LiCre and LiCre-V5 following pulsed photo-stimulation. Cells were subjected to pulses of blue light (450 nm, period of 10 s, duty cycle of 10%, intensity of 35 mW/cm2) for 1 hour, then transferred to fresh medium for 4 hours to enable expression of the fluorescent proteins. The fraction of cells expressing the mCherry reporter (ON cells) was determined by flow cytometry. *P*-value: Welch *t*-test. **d)** Recombination rate induced by LiCre in conditions of riboflavin supplementation. Same illumination conditions as in c). Medium was supplemented with various concentrations of riboflavin (x-axis). Blue, LiCre expressed under the Pmet17 promoter (plasmid pGY466). Red, LiCre expressed under the Pmet17 promoter and FMN1 expressed under the Padh1 promoter (plasmid pGY781). Dots: independent cultures (biological replicates). Lines and ribbons: mean +/− s.e.m.. **e)** Recombination rate following pulsed stimulation of various duration. Cells were subjected to pulses of blue light (450 nm, period of 10 s, duty cycle of 5%, intensity of 35 mW/cm2) for the indicated amount of time, then transferred to fresh medium for 4 hours to enable expression of the fluorescent proteins. Green, red and non-fluorescent cells were then counted by flow cytometry. **f**) Same as in *e* but for a single duration and various temperatures. **g**) Same as in *e* but with various periods and a duty cycle of 10%. Intensity of 35 mW/cm2. **h**) Same as in *e* and *g* but with various light intensities, a period of 7 s, a duty cycle of 29 %, applied for 60 minutes. Colors correspond to two independent sets of experiments. In *e-h*, replicates (dots) correspond to independent cultures, each inoculated with an independent transformant of the LiCre plasmid. Lines connect mean values.

We first used the reporter yeast strain to compare the efficiency of LiCre with its epitope-tagged version LiCre-V5. After one hour of photo-stimulation (1s-pulses of light applied every 10 seconds), the fraction of cells that had recombined the mCherry reporter (ON cells) was ∼40% in the case of LiCre, and only ∼20% in the case of LiCre-V5 (Fig. 4c). This difference in efficiency was statistically significant (p-value = 0.013, Welch t-test); it showed that the presence of repeated V5 epitopes at the C-terminal end of LiCre reduces its activity. It is possible that this tag negatively affects the DNA-binding affinity of LiCre, which would explain the relatively low ChIP signal observed above for LiCre-V5 *in vivo* (Fig. 1) as compared to the strong DNA-binding affinity measured in vitro for LiCre (Figs. 2 & 3).

We then evaluated whether the photo-activation of LiCre could be limited by the availability of intra-cellular flavin mononucleotide (FMN). FMN is an essential co-factor for the photo-induced conformational switch of LiCre’s LOV domain. Cells produce it by phosphorylation of its precursor riboflavin. To test if LiCre efficiency would increase in conditions that maximize FMN production, we applied our assay on cells cultured in the presence of various concentrations of riboflavin (up to 20 µM). As shown in Fig. 4d, the presence of extracellular riboflavin had no significant effect on recombination efficiency. To exclude the possibility that riboflavin was insufficiently processed into FMN, we added in the LiCre-encoding plasmid a cassette encoding the over-expression of the yeast riboflavin kinase Fmn1p (constitutive expression conferred by the Padh1 promoter). In this case, a modest increase of LiCre efficiency was seen at 2 µM and 5 µM riboflavin, but not at higher concentrations (10 and 20 µM) (Fig. 4d). We conclude that, in standard conditions, FMN is probably sufficiently abundant in yeast cells for the optimal activation of LiCre.

Figure 4e shows how recombination efficiency increased with the duration of a defined pulsed stimulation. In agreement with previous observations [1], ten minutes were enough to induce recombination in a significant proportion (∼10%) of cells. In general, we counted slightly more red cells than green cells. This could indicate that the reporter locus on chromosome IV is more favorable to recombination than its counterpart on chromosome II. Alternatively, the difference could result from different measurement accuracy between the green and the red channel. Importantly, we never observed recombination in the entire cell population. After prolonged (>3 hours) photo-stimulation, the proportion of positive cells reached a plateau of about 85%. Most of these cells displayed both red and green fluorescence, leaving a remaining ∼15% cells negative for both reporters. This suggested that the upper limit of recombination efficiency was caused by a cell-specific factor rather than a locus-specific factor. We hypothesized that this upper limit could be linked to the occasional loss of the LiCre-encoding plasmid.

To test this possibility, we repeated the experiment but instead of illuminating the population of cells, we plated diluted samples of it on both selective and non-selective solid media (Supplementary Table S6). This revealed that, in our experimental conditions, the proportion of cells having lost the LiCre plasmid at the time of illumination was between 10% and 30%. This observation is remarkably consistent with the presence of ∼15% of recombination-negative cells in our optogenetic assay with prolonged photostimulation. We therefore propose that, for cells having kept the plasmid, LiCre-mediated recombination efficiency after prolonged illumination is close to 100%.

We also observed substantial variability between experiments. For example, a pulsed stimulation of 60 minutes generated nearly 50% of positive cells in the experiment of Fig. 4e, whereas the same stimulation generated only ∼30% of positive cells in another experiment (see Fig. 3g of [1]). We suspected that variations in room temperature could contribute to this variability and we tested this possibility by quantifying recombination efficiency in various thermo-controlled conditions. The same pulsed stimulation generated between ∼15% and ∼45% positive cells when temperature varied from 20°C to 37°C, respectively (Fig. 4f), higher temperature leading to more efficient recombination, thus explaining most of the variability mentioned above. We therefore conducted subsequent experiments under strict temperature-controlled conditions (30°C).

To further explore the dynamic properties of the LiCre-*loxP* reaction *in vivo*, we applied pulsed photo-stimulation at various periods, with two different duty cycles (fraction of the period when light is on). The list of experiments is summarized in Supplementary Table S5 and only a subset of the data is shown in Fig. 4g. For a fixed total cumulative time of light exposure, recombination efficiency declined when the period increased. For example, applying a total illumination of 6 minutes (dark red in Fig. 4g) led to an efficiency above 40% when the time between consecutive pulses of light was short (4.5 seconds) but below 25% when this period was longer (54 seconds). This may reflect the existence of a lag time, in the order of tens of seconds, before LiCre returns to its dark, inactive, state in absence of light.

In order to quantify the photo-sensitivity of the reaction, we illuminated yeast cells at various light intensities. We chose a regime where light pulses were long enough to ensure efficient activation (2 seconds) and frequent enough to prevent the system from reverting to its inactive state (every 7 seconds). As expected, we observed an overall increase of recombination efficiency with light intensity until maximal activation was reached at 12mW/cm^2^ (Fig. 4h, red). Intriguingly, the fraction of positive cells was slightly lower when we applied maximal intensity (35.6mW/cm2). This observation was reproduced when we repeated the experiment (Fig. 4h, black, Dataset 11 listed in Supplementary Table S5), possibly pointing to photo-toxic effects under prolonged or intense illumination.

Overall, the experimental data generated here describe the efficiency of LiCre following illumination conditions varying in duration, in dynamics or in light intensity. This dataset associates the kinetic properties of the input stimulus with the quantitative output of the system.

### Kinetic modeling of LiCre-loxP recombination

To describe the light-induced recombination catalyzed by LiCre, we built a mathematical model based on three steps (Fig. 5, see Methods).

**Figure 5.**
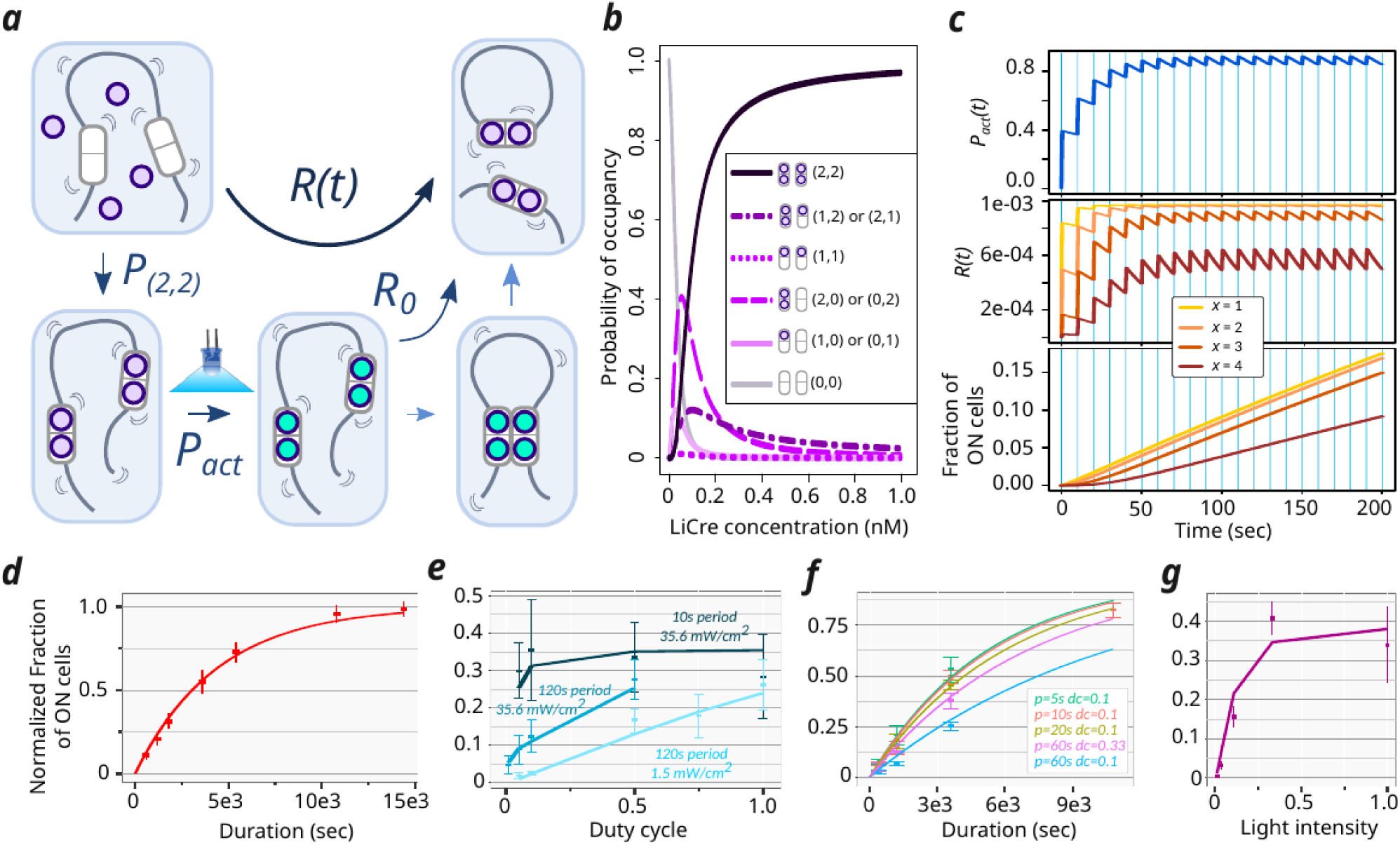
Kinetic model of LiCre-*loxP* recombination. **a)** Model description. Purple and cyan beads represent inactive and active LiCre units, respectively. Occurrence of the reaction assumes the binding of four LiCre units (which happens with probability P_(2,2)_), the photoactivation of at least *x* of them, with *x* ɛ {1,2,3,4}, and a maximal reaction rate *R_0_*once the complex can form. Reaction rate *R(t)* summarizes the triggering of the reaction by light. See text and Methods for details. **b)** Effect of [LiCre] concentration on the probability of occupancy at two *loxP* sites. (*n,m*) indicates *n* and *m* LiCre molecules bound to the first and second *loxP* site, respectively. **c)** Simulations under periodic pulses of light. Light was ON during the vertical blue stripes, each pulse occurred every 10 s and lasted 0.5 s. [LiCre] = 1 nM, *k_ON_* = 1 s^−1^, *k_OFF_* = 0.007 s^−1^, *R_0_* = 10^−3^ s^−1^. From up to bottom: probability that one LiCre unit is active (*P_act_*); reaction rate at each possible value of *x* (required number of activated units); cumulative fraction of recombined molecules over time at each possible value of *x.* **d-g)** Best model prediction using *x* = 2 and parameters simultaneously fitted to all experimental data. Lines, model simulations. Dots and bars, experimental data (mean ± 2 standard deviations). Each panel corresponds to one consistent series of acquisitions, where the fraction of mCherry-positive yeast cells which underwent recombination was measured by flow-cytometry (datasets 1-10 listed in Supplementary Table S5). Value *k_OFF_* inferred from all data: 50 s^−1^. **d)** Effect of the duration of illumination (same data as in Fig. 4e). Inferred *R_0_* value: 3.5.10^−4^ s^−1^. **e)** Effect of illumination dynamics and intensity (duration of 1 hour), data from Fig. 3f-g of ref. [1]. Inferred *R_0_* value: 0.000125 s^−1^. **f)** Effect of illumination dynamics and duration. Inferred *R_0_*value: 0.00014 s^−1^. **g)** Effect of illumination intensity (same data as in Fig. 4h red). One unit = 35.6 mW/cm^2^. Inferred *R_0_* value: 0.00022 s^−1^.

First, association of LiCre to DNA occurs in the dark and is at equilibrium. Using the dissociation constants inferred from the SPR experiments (see above), we plotted the probabilities of occupancy of the four LiCre binding sites located on two *loxP* sequences assuming independent binding at the two *loxP* loci (Fig. 5). At LiCre concentrations above 1nM, over 97% of cells have both their *loxP* sites fully occupied (four bound LiCre units in total).

Second, during stimulation by light, bound LiCre units are activated at a rate *k_ON_*. In dark condition, they deactivate at a rate *k_OFF_*. Given these rates and the dynamics of illumination, we can compute the probability *P_act_(t)* that a given LiCre unit is active at time *t* (Fig. 5c, top). Since LiCre is activated by the asLOV2 switch, we chose *k_ON_* as previously reported for the LOV2 domain [13]. This rate increases linearly with the incoming flux of photons and is in the order of second^−1^ (see Methods).

Third, if the two *loxP* sites are both fully-occupied and if at least *x* among the four LiCre units are active, with *x* ranging from 1 to 4, recombination can occur at a constant rate *R_0_* that encompasses the loop probability between the sites, the synapse formation and the actual catalytic reaction of recombination. All this leads to an overall time-dependent recombination rate *R(t)* that is proportional to *R_0_*, to the probability of having 4 LiCre bound to the *loxP* sites and to the probability to have at least *x* light-activated LiCre molecules, which is a function of *P_act_(t)* (Fig. 5a).

The well-known Cre-*loxP* reaction is reversible and this is presumably also the case for the LiCre-*loxP* reaction. As described above, we concluded from our experimental data that all cells having kept the LiCre plasmid would eventually achieve recombination after prolonged photo-activation. Thus, at equilibrium, the backward reaction (re-insertion of the excised circular DNA) is unlikely and out-competed by the forward reaction and we therefore neglected it in our model.

A reaction simulated with this model is shown in Fig. 5c. In this example, periodic pulses of light were applied and the fraction of activated LiCre units reached a periodic steady-state after only 60 seconds, oscillating around ∼80% (Fig. 5c, top) with the light stimulus. The overall time-dependent reaction rate *R(t)* also fluctuates, the speed of recombination being highly affected by the number (*x*) of active LiCre units needed to form a functional complex (Fig. 5c, center). After plotting the cumulative fraction of recombined molecules over time (Fig. 5c bottom) we concluded that this three-steps model provides realistic outcomes that can be compared to experimental data.

### Model fitting to experimental data suggests that photo-activation of two LiCre protomers may be sufficient to trigger recombination

We investigated if the above model could explain the efficiencies of DNA recombination that we observed on yeast cells stimulated under various conditions. In this yeast assay, the fraction of cells that switched ON fluorescence directly reports on the fraction of DNA molecules (among the total cell population) that underwent recombination. The corresponding experimental data can therefore be compared with model predictions.

Although we did not quantify LiCre concentration in yeast cells, we estimated from the literature (see Methods) that it was probably above ∼1nM, conferring near-complete occupancy at the two *loxP* sites *in vitro* (Fig. 5b). Arbitrarily, we decided to set LiCre concentration in the model at 1nM. Note that this choice does not impact the quality of the fit nor the order of magnitudes of the inferred parameters (see Methods).

In total, three parameters of the model are unknown and can be optimized to fit experimental data: *k_OFF_*, the speed of LiCre inactivation in the dark; *x,* the minimal number of active LiCre protomers in the complex required for the recombination reaction to occur; and *R_0_*, the reaction rate of the enabled synapse. We ran the optimization of *k_OFF_* and *R_0_* for each value of *x* among {1, 2, 3, 4} over a rich set of experiments performed under various conditions of illumination (different durations, periods, duty cycles and light intensities, data sets 1-10 listed in Supplementary Table S5). Given the batch-to-batch experimental variability mentioned above, we optimized *R_0_* specifically for each series of experiments. Finally, to account for the frequent loss of the LiCre plasmid (see above), we normalized experimental data so that the highest observed efficiency was 100% (see Methods).

Results from parameter optimization are summarized in Table 1. Inferred *R_0_* values ranged from 1.25 to 4.10^−4^ s^−1^ (characteristic time from 42 min to 2.2 hrs) independently of *x* and this variation primarily corrected for the above-mentioned suspected effect of temperature in one of the data sets. When *x* was set to either 1, 2, 3 or 4, the inferred *k_OFF_* rates corresponded to characteristic LiCre deactivation times of about 8s (*k_OFF_* = 125.6 × 10^−3^ s^−1^), 20s (*k_OFF_* = 50 × 10^−3^ s^−1^), 50s (*k_OFF_* = 19.9 × 10^−3^ s^−1^) and 140s (*k_OFF_* = 7.1 × 10^−3^ s^−1^), respectively. It has long been known that, once photo-activated, the LOV2 domain needs tens of seconds to return to its dark state [14]. It is therefore unlikely that LiCre can be deactivated after only 8 seconds in the dark as predicted by the model where a functional recombination complex is formed if only one LiCre unit is active (*x* = 1). Notably, adequacy of model predictions to the data was maximized for *x* = 2 (Table 1), with an overall very good fit (Fig. 5d-g), and using *x* = 4 provided the least adequate match to experiments but still capturing well most of the data (Table 1). This suggests that the complex may not need four active LiCre units to be functional, but that photo-activation of only two units may be sufficient.

**Table 1:**
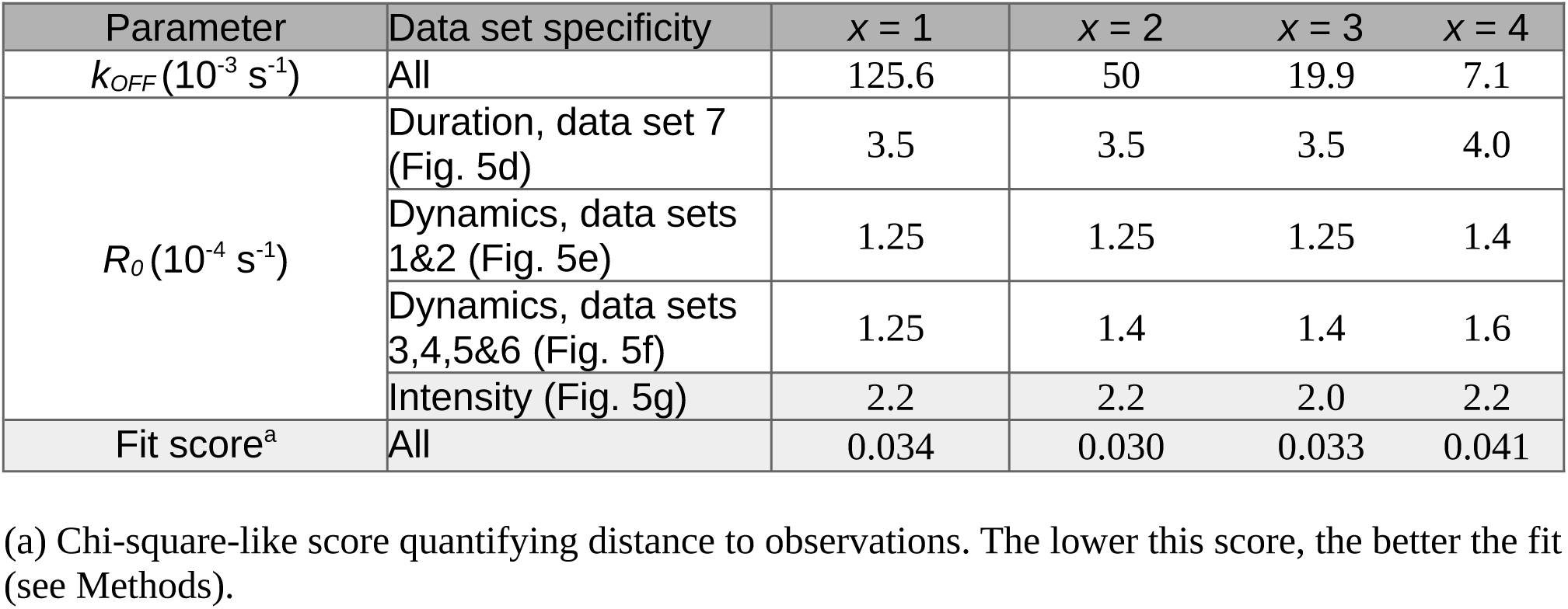
Best-fit parameters of the recombination reaction model.

| Parameter | Data set specificity | $x = 1$ | $x = 2$ | $x = 3$ | $x = 4$ |
| --- | --- | --- | --- | --- | --- |
| $k_{OFF}(10^{-3} \text{ s}^{-1})$ | All | 125.6 | 50 | 19.9 | 7.1 |
| $R_0(10^{-4} \text{ s}^{-1})$ | Duration, data set 7 (Fig. 5d) | 3.5 | 3.5 | 3.5 | 4.0 |
|  | Dynamics, data sets 1&2 (Fig. 5e) | 1.25 | 1.25 | 1.25 | 1.4 |
|  | Dynamics, data sets 3,4,5&6 (Fig. 5f) | 1.25 | 1.4 | 1.4 | 1.6 |
|  | Intensity (Fig. 5g) | 2.2 | 2.2 | 2.0 | 2.2 |
| Fit score<sup>a</sup> | All | 0.034 | 0.030 | 0.033 | 0.041 |

(a) Chi-square-like score quantifying distance to observations. The lower this score, the better the fit (see Methods).

### The model captures the effect of a point mutation affecting the light cycle of LiCre

We tested if the response of LiCre to pulsed stimulation was modified by mutations altering the light-cycle of its asLOV2 domain. Several point mutations were previously described that either slow down or accelerate the recovery of asLOV2 in the dark [15–18]. Using earlier reports, we selected 3 mutants with faster recovery (V416T, N425Q/I427V and T418S)[18] and 3 mutants with slower recovery (V416L, F494I/L496S, and Q513L)[15,18]. We introduced these mutations in LiCre and, using our yeast assay, we quantified the recombination efficiency of the resulting 6 variants under a given regime of periodic photo-stimulation. All mutations severely affected the performance of LiCre (Fig. 6a). Photo-activation was lost in the slow-recovery mutant Q513L. Four mutants (V416L, F494I/L496S, V416T and N425Q/I427V) showed a pronounced residual activity in the dark. Photo-activation was preserved in the remaining mutant, T418S, but it was reduced as compared to wild-type LiCre.

**Figure 6.**
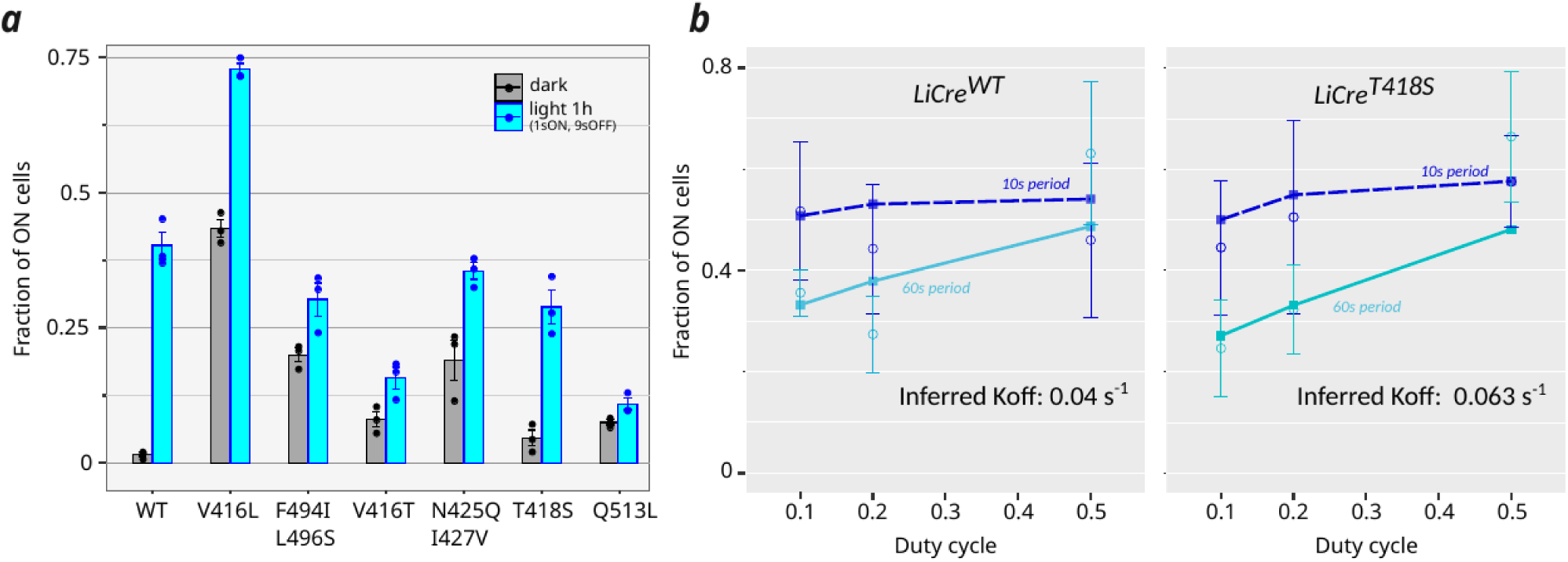
Performance of LiCre mutants with altered photo-cycle of the LOV domain. **a)** Recombination induced by wild-type (WT) LiCre and 6 genetic LiCre variants. Efficiency (fraction of ON cells) was quantified by flow-cytometry following pulsed photo-stimulation (450 nm, period of 10 s, duty cycle of 10 %, intensity of 35 mW/cm2, duration of 1 hour). **b)** Recombination efficiencies of LiCre WT and T418S variant following various regimes of pulsed photo-stimulation (Dataset 13 of Supplementary Table S5). Circle dots: mean values of experimental data for a duration of 60 min. Error bars: +/− standard deviation of experimental data. Squares connected by lines: predictions from model with *x*=2 and best-fit parameters *R_0_* and *k_OFF_* (Supplementary Table S7).

We reasoned that if this reduction resulted from a faster recovery of LiCre to its inactive state, then increasing the duration or the frequency of the light pulses would restore activity in this mutant, possibly up to wild-type levels. To test this prediction, we quantified the recombination efficiencies of both LiCre^WT^ and LiCre^T418S^ following photo-stimulation regimes for two different periods and three different duty cycles (Fig. 6b). As expected, for a fixed duty-cycle, the recombination efficiency of both constructs was higher under a short (10s) period than under a long (60s) period; and similarly, for a fixed period, it increased with the duty-cycle, especially for the 60s period. The main difference between the two variants was seen at 10% duty-cycle and a period of 60s. Under this condition, the mean fraction of switched cells was 0.355 for wild-type LiCre, and 0.246 for the T418S mutant, a difference that was statistically significant (*p*-value = 0.005, Welch *t*-test). At 50% duty-cycle, and for both periods, the two constructs displayed similar efficiencies. Thus, the effect of the T418S mutation manifests only in sub-optimal regimes where the time for inactivation in the dark is long. This observation is consistent with the prediction that the T418S mutation increases the recovery rate of LiCre.

Given this consistency, we evaluated if our kinetic model was able to capture such a difference in recovery rate between the two constructs. By fitting the comparative experiments described above (Dataset 13 listed in Supplementary Table S5) with our kinetic model, we inferred *k_OFF_* and *R_0_* parameters for LiCre^WT^ and LiCre^T418S^ independently. We did so separately for all four *x* values (=1, 2, 3 or 4, the required number of active units). Corresponding model predictions for *x*=2 are shown in Fig. 6b, but all parameter estimates are provided in Supplementary Table S7. All four models inferred *k_OFF_* values that were higher for the mutant than for the wild-type. This difference increased from 1.58-fold to 1.99-fold when *x* increased from 2 to 4. This trend is consistent with the fact that a model imposing that all four units must be active for the reaction (*x*=4) is more sensitive to the inactivation rate (*k_OFF_*) of individual units. From the different estimates of *k_OFF_* between the LiCre variants, we conclude that the faster asLOV2 recovery caused by the T418S mutation translates into a ∼1.6-fold faster inactivation of LiCre in the dark.

In conclusion, our three-steps model can recapitulate the recombination rates observed under various dynamics and intensities of light stimulation; it suggests that synapse assembly may occur even if not all four LiCre units are photo-activated; it provides quantitative estimates of the reaction rate of the assembled synapse and of the speed of monomer’s recovery to the dark state; and it captures an expected faster recovery resulting from the T418S point mutation of the LOV domain of LiCre.

## DISCUSSION

By coupling modeling with experiments conducted both *in vitro* and *in vivo*, this work provides knowledge on the kinetics of site-specific DNA recombination by blue-light photo-stimulation of the LiCre-*loxP* system. We found that LiCre has high affinity for its target *loxP* substrate even if it is not photo-activated, that this binding is cooperative – although less than for Cre - and that the observed data are best explained by a model where the photo-activation of at least two LiCre protomers enables recombination. We quantified LiCre DNA-binding affinity and we estimated characteristic times involved in the photo-induced reaction. We identified a point mutation altering the light cycle of LiCre.

The fact that LiCre binds its DNA target already before illumination probably explains why it is faster and more efficient than other photo-activatable Cre-*loxP* systems [1]. These other systems were obtained by splitting Cre in two halves which were fused to optogenetic dimerizers enabling their assembly in response to light [4–7]. Given that Cre binds DNA by capturing it between its N- and C-terminal domains [8], it is unlikely that these split subunits can bind DNA before they are assembled together. In contrast, LiCre is made of a single peptide chain comprising both N- and C-terminal domains, which makes it compatible with DNA binding prior to illumination. In addition, the single-chain design of LiCre, together with its high affinity for its target DNA, predict that it may enable efficient recombination over a wider range of intracellular concentrations than split-based designs. This advantage may be important when intracellular concentration of the enzyme is unknown or difficult to control.

Our SPR measurements allowed us to compare the DNA-binding affinities of LiCre and Cre. For Cre, we estimated K_D1_ at 1.15 nM, which is concordant with previous reports where it was estimated at 1.7 nM by electrophoretic mobility shift assays (EMSA) [12] and at 1.8 nM by SPR [11]. For K_D2_, however, these previous studies reported values of 0.0135 nM for EMSA and 0.02 nM for SPR, which are much higher than 0.001 nM found in the present work. This difference, which results in different estimates of DNA-binding cooperativity, can be explained by differences in Mg^2+^ concentration between the different studies. Unlike previous reaction conditions which did not contain any Mg^2+^ cations [11] or contained 2 mM Mg^2+^ partially chelated by 1 mM EDTA [12], our running buffer contained 10 mM MgCl_2_ and no EDTA. Magnesium as well as spermidine have long been known to increase the activity of Cre *in vitro* [19], and the previous SPR study reported a marked increase in cooperativity when one of these ingredients was added to the reaction. For example, the DNA-binding cooperativity moment of Cre increased from 86 to 1700 when 5 mM spermidine was added, and the authors reported that the effect of Mg^2+^ was similar [11]. Thus, the high cooperativity previously observed in the presence of Mg^2+^ is comparable to the one we found here (cooperativity moment of 1300). We conclude that our measure of the DNA-binding affinity of Cre by SPR is consistent with previous estimates, which provides confidence in the novel measures obtained here on LiCre.

Similarly to Cre, LiCre showed very high affinity for *loxP,* and LiCre proteins bind cooperatively to DNA. However, when compared to Cre, we observed a 2-fold stronger binding of the first unit to DNA and a 7-fold weaker binding of the second unit, resulting in a much less pronounced cooperativity (Fig. 3f). Given that cooperativity results from physical interactions between protomers, this observation is consistent with the way LiCre was engineered. Indeed, the original protein-protein interactions known to take place between Cre protomers were destabilized in three ways [1]. First, the αA helix of the N-terminal domain was truncated. Second, two point mutations destabilizing protein-protein interactions were introduced in the αN helix of the C-terminal domain. These point mutations likely affect the strength of the mechanism by which this helix was recently shown to participate to the DNA-binding cooperativity of Cre: in the absence of DNA, soluble Cre is monomeric and retains its own αN helix in *cis*, but in the presence of *loxP* DNA, the αN helix is released and becomes an anchor that stabilizes the association of the second unit [20]. This release of helix αN probably also occurs for LiCre, but likely with a lower contribution to cooperativity because the two mutations make αN a weaker anchor. Finally, the remaining segment of the N-terminal αA helix of Cre was directly fused to the Jα helix of asLOV2. Thus, the LOV domain of LiCre is located in its N-terminal globule, away from the C-terminal αN helix, and the stabilization mediated by the C-terminal module is probably not light-dependent; in agreement with the remaining cooperativity of LiCre observed here in the dark. Photoactivation of LiCre is expected to result from conformational changes occuring at its N-terminal: in the dark, the chimeric Jα-αA helix is probably unable to interact with the partner protomer because it is presumably well-folded and sequestered by the asLOV2 domain; illumination is expected to cause the release of this helix, allowing it to refold and participate to inter-unit interaction. Given the multiple changes that destabilize protein-protein interactions, it is not surprising to observe a weaker DNA-binding cooperativity for LiCre than for Cre. It would be interesting to measure the DNA-binding affinity of light-activated LiCre. We may expect an increase in cooperativity if the chimeric helix refolds in a conformation suitable for protein-protein interaction, but probably not to the extent of Cre given the C-terminal mutations. Such measurements require dedicated equipment that can simultaneously quantify molecular associations and illuminate samples in appropriate conditions, which was not possible on our SPR station.

Unlike classical genetics or drug-based treatments, optogenetics offers the possibility to induce an intracellular reaction dynamically with high temporal precision. In the present study, applying pulsed stimuli with various dynamics and feeding the resulting data into a simple kinetic model allowed us to derive kinetic information from a rather simple yeast-based assay. We could conclude that, following their photo-activation, LiCre units get deactivated in tens of seconds, that the LOV-targeting point mutation T418S accelerates this de-activation, that a functional LiCre-*loxP* synapse may be able to form if at least two of its four LiCre units are active and that, following this activation, it achieves DNA recombination quite slowly with a characteristic time of tens to hundreds of minutes. It is interesting to compare these conclusions with the kinetic properties of Cre-mediated recombination, which were previously studied by tracking the motion of single tethered DNA molecules *in vitro* [21–23]. These studies showed that, with a constitutively-active Cre protein, limiting steps of the reaction were the assembly of a functional synapse and the catalytic activity of the complex. For example, for intra-molecular recombination reactions between *loxP* sites separated by 3044 and 870 bp, transposing the estimates of reaction rates found in [23] to equivalents of the *R_0_* parameter of our model (see Methods) leads to values of 0.2-0.55 10^−4^ s^−1^, respectively, corresponding to characteristic times reaching several hundreds of minutes, much slower than our *in vivo* estimation for two *loxP* sites separated by 1300 bp. Since, for LiCre, the steps of synapse assembly and catalysis are not expected to be faster (LiCre was derived from Cre by reducing, rather than increasing, its propensity to assemble the complex and achieve the reaction), the main difference between *in vitro* [23] and *in vivo* (this study) R_0_ values likely lies into the probability of forming a DNA loop bringing the two *loxP* sites together (a pre-requisite to synapse formation). This probability depends on the mechanical properties of the inter-*loxP* DNA segment [24,25]; it may thus differ strongly between a stiff inter-*loxP* segment of naked dsDNA *in vitro* and a softer, chromatinized one *in vivo* [26,27]. Nonetheless, we note that some of our conclusions remain to be validated by additional direct mechanistic investigations. Single-molecule analyses, such as those previously performed on Cre [21,28], would enable to evaluate more complex models. For example, it is possible that the reaction rate of a fully-activated synapse may be faster than the rate of a synapse containing only two or three active LiCre units. Also, *in-vitro* kinetic data might better discriminate models with different *x* values. Complex assembly after activation of two units is consistent with the need of two contact interfaces to bridge the two dimer-*loxP* complexes together, however this prediction was supported here by relatively mild differences in fit scores to the yeast-based data (Table 1).

Remarkably, the model was able to capture a quantitative change in the inactivation rate (*k_OFF_*) of LiCre resulting from a point mutation targeting its LOV domain (Fig. 6b). This mutation, referenced T418S on the Phototropin1 sequence, was initially isolated from a screen where asLOV2 recovery was tracked by monitoring the fluorescence of bacterial colonies [18]. The recovery rate of this mutant was reported to be ∼14 times faster than the recovery rate of wild-type asLOV2. Here, we find that introducing this mutation in LiCre increases its inactivation rate to about 1.6-fold. It is important to note that the rates of asLOV2 recovery and the rate of LiCre inactivation do not describe the exact same biochemical steps. The former corresponds to the conformational change of the LOV domain from a photo-stimulated (light) to its original (dark) state; the latter corresponds to the loss of activity of the full LiCre protomer, which involves not only the conformational change of its LOV domain but also the resulting putative loss of functional interaction with other LiCre protomers of the recombination synapse. It is therefore not surprising to observe a milder effect of the mutation on the speed of LiCre de-activation than on the speed of asLOV2 recovery.

Finally, our results have practical implications on experimental protocols employing LiCre. First, given its high affinity for *loxP* (Fig. 5b), over-expressing LiCre at high levels will probably not increase its efficiency. On the contrary, depending on how cells regulate the production of the flavin mono-nucleotide (FMN), an overproduction of LiCre might generate a proportion of LiCre molecules lacking this co-factor essential for photo-activation. Here, in the conditions used in our yeast-based assay, FMN is probably abundant enough in the cell to occupy all LiCre molecules. Otherwise, adding riboflavin to the culture medium and over-expressing the riboflavin kinase would have generated a higher fraction of switched cells in the assay; which was not observed (Fig. 4d). In mammalian cells, FMN was previously quantified at about 1 amol per cultured cell [29] (equivalent to 600,000 molecules, or 1 µM assuming a cell volume of 1 pL). Thus, it may be judicious to express LiCre at moderate levels (< 1 µM) to ensure stoechiometry with FMN while still maximizing its binding to *loxP*. However, it is also possible that, if high levels of LiCre would cause a sequestration of FMN from the cellular pool, then cells might be able to respond by increasing FMN synthesis. This would provide enough co-factor for most LiCre molecules regardless of the expression level of LiCre.

Second, LiCre was three times more efficient at 37°C than at 20°C, with no additional residual activity in the dark (Fig. 4f). This observation is concordant with previous *in vitro* measurements for Cre [30]. We therefore invite LiCre users to conduct experiments at 37°C whenever possible.

Third, our data indicate how to minimize the amount of blue light shone on cells – and its potential damaging effects thereof - while keeping LiCre induction maximal. With a 450 nm wavelength, we recommend to use a photon flux of 12 mW/cm^2^ and periodic pulses of 1 second spaced every 10 seconds.

In conclusion, by precisely characterizing the kinetic properties of the optogenetic LiCre-*loxP* recombination, this study provides useful insights for conducting targeted genetic manipulations of live cells using light.

## MATERIALS AND METHODS

### Strains and plasmids

Plasmids, strains and oligonucleotides used in this study are listed in Supplementary Tables S1-3. Records of laboratory stocks were traced using MyLabStocks [31].

To perform chromatin immuno-precipitation (ChIP), we tagged LiCre with V5 epitopes (IPNPLLGLD) [32]. We placed a Gly-Ser flexible linker at the C-terminus of LiCre in order to avoid interference between these epitopes and the C-terminal αN helix known to be critical for LiCre activity [1]. Nine repeats of the V5 epitope were amplified, using primers 1Q46 and 1Q47, from a plasmid kindly provided by Pascal Bernard. The resulting amplicon was co-transformed in yeast with plasmid pGY466 previously linearized by NdeI. The plasmid (pGY605) resulting from subsequent homologous recombination was recovered from yeast colonies and amplified in *E. coli*. We constructed Cre-V5 similarly by co-transforming in yeast the same amplicon with pGY502 linearized at NdeI. Subsequent rescue from yeast provided plasmid pGY607. Both plasmids were verified by Sanger sequencing, which revealed that pGY605 contained nine V5 repeats as expected and pGY607 unexpectedly contained twelve V5 repeats.

For recombinant protein production, the LiCre and Cre coding sequences were codon-optimized for *E. coli* and the corresponding sequences were synthesized and cloned in pET-19 by Genecust to produce plasmids pGY611 (pET-LiCre) and pGY709 (pET-Cre), respectively. An mCherry reporter targeting the *LYS2* locus was obtained by cloning a KpnI-AvrII fragment from pGY472 into the KpnI-AvrII sites of pGY537, producing plasmid pGY618. For integration of a single *loxP* site at the *LYS2* locus, we built plasmid pGY621 by digesting pGY618 with BamHI and AgeI, and applying Klenow fill-in and re-ligation. LiCre mutants targeting the asLOV2 domain were generated by Genecust via mutagenesis of plasmid pGY466. Note that positions V416, F494, L496, N425, I427, T418 and Q513 referenced on Phototro pin1 correspond to physical positions V25, F103, L105, N34, I36, T27, and Q122, respectively, on the amino-acid sequence of LiCre. An expression cassette consisting of the promoter of the *S. cerevisiae* ADH1 gene, the coding sequence of *S. cerevisiae* FMN1, and the TEF terminator sequence from *Ashbya gossypii* was synthesized by Genecust who cloned it into the KpnI site of pGY466 to produce pGY781.

Reporter yeast strains were either GY2214 [1] carrying an mCherry reporter at the *HO* locus and a GFP reporter at the *LYS2* locus, or strain GY2517 carrying the same mCherry and GFP reporters, but at the *LYS2* and *HO* locus, respectively, or strain GY2753 carrying only the mCherry reporter at the *HO* locus. To obtain GY2517, we transformed GY855 with pGY618 linearized by NruI and selected Lys-, Ura+ clones (pop-in) and then 5-FoA resistant clones that had become Lys-, Ura- (pop-out). This produced strain GY2450 which we crossed with GY1761 to obtain GY2517. To obtain GY2753, we first cured the ade2-1 mutation of strain OAy470 [33] by transforming it with a PCR amplicon containing the wild-type ADE2 sequence. The resulting strain (GY2752) was then transformed with a 4Kb NotI fragment from pGY472.

For ChIP, strain GY2416 containing a single *loxP* site at the *LYS2* locus on the right arm of chromosome 2 was obtained by transforming strain GY855 with plasmid pGY621 linearized at NruI followed by pop-in and pop-out selections as above.

### Purification of recombinant Cre and LiCre from bacteria

The coding sequences of Cre and LiCre fused to a N-terminal 6xHIS tag were codon-optimized for production in *E. coli*. They were synthesized by GeneCust and cloned in a pET-19 vector. The resulting plasmids (pGY611 and pGY709) were transformed in Rosetta2 bacteria. Bacterial cells were cultured in LB medium containing ampicillin and chloramphenicol at 37°C. A 40 mL starter culture was used to inoculate a 1 L culture. When reaching OD_600_ = 0.6, these cultures were induced for 3 h with 0.1 mM IPTG. Cells were harvested by centrifugation and resuspended in 50 mL of 50 mM Tris pH8, 500 mM NaCl, 10% glycerol, 0.1% Triton, 0.5 M glucose, 1 mM DTT, 1 g/L lysosyme, 2X mix of protease inhibitors, 5 µg/mL DNAse and 2.5 mM MgCl2. After 30 minutes on ice, the suspension was sonicated 4 times for 1 min and then centrifugated at 14,000 g for 20 min at 4°C. The resulting supernatant was filtered on a GD/X membrane (INI) and then loaded on a HisTrap 1 mL column (GEHealthcare) using an AKTA PURE chromatography system and a varying ratio of two buffers A and B. Buffer A contained 50 mM Tris pH8, 500 mM NaCl, 10% Glycerol. Buffer B had the same composition of A with imidazole added at 300 mM. Three series of injections and washes were performed using A:B mixtures of different proportions : (100% : 0%), (95% : 5%) and (90% : 10%). Elution was done by applying a gradient of A:B mixtures from (90% : 10%) to (50% : 50%) which lasted 10 CV (column volume taking into account the packing and porosity of the material), followed by a (0% : 100%) mixture for 5 CV. Fractions were analyzed by SDS-PAGE. We noted that the eluted fractions of interest had a yellow color (Fig. 2a), indicating that LiCre was co-purified with its co-factor flavin mononucleotide (FMN). Fractions of interest were concentrated on a 10 kDa membrane (Vivaspin, Millipore). The resulting products were purified through two columns: a GF Superdex75 10/300 Increase column (Cytiva) and a Zeba column which was eluted in the final SPR buffer 20 mM Tris pH7.4, KCl 50 mM, NaCl 150 mM, MgCl_2_ 10 mM, DTT 1 mM, P20 surfactant 0.05%. The quality of the production was assessed by light diffusion on a NanosizerS (MALVERN) and SDS-PAGE (Fig. 2b).

### Chromatin immunoprecipitation (ChIP)

Strain GY2416 was transformed with plasmid pGY44, pGY605 or pGY607. Three independent transformants were used as biological replicates (9 strains in total). For each replicate, the strain was cultured overnight in 120 mL liquid flasks of synthetic SD-MW medium containing 2% glucose. Composition of this medium was the same as for the SD-Met medium previously described [34], except that tryptophan was missing in the Dropout Mix. The next day, 54 mL of each culture was loaded on a 6-well plate (9 mL per well) and illuminated in a DMX-controlled 450 nm-LED box placed in a 30°C incubator. Light intensity was set to 35 mW/cm^2^. Illumination consisted of periodic pulses of 1 s applied every 10 s for a total duration of 60 minutes. In parallel, 54 mL of the same culture was kept in the dark. For each sample (illuminated or not), 25 mL was processed for Western Blot (WB) and 25 mL was processed for ChIP. For WB, cells were washed twice with TBS1X, aliquoted in microtubes at 10^8^ cells per tube and pelleted by centrifugation. For ChIP, 700 µL of 37% Formaldehyde (Sigma F8775) was added to the 25 mL cell suspension. After 15 min at room temperature, 1250 µL of 5M Glycine was added and tubes were placed on a rotating wheel for 5 min at room temperature. Cells were then washed twice with TBS1X, aliquoted in microtubes at 10^8^ cells per tube and pelleted by centrifugation. All cell pellets (WB and ChIP) were frozen in liquid nitrogen and stored at −80°C.

For WB, cell pellets were thawed on ice and resuspended in 300 µL lysis buffer composed of 0.1% w/v sodium deoxycholate, 1 mM EDTA pH8, 50 mM HEPES-KOH pH7.5, 140 mM NaCl, 1% w/v Triton X-100 and proteinase inhibitors (Sigma P2815), mixed by pipeting and transferred to Sarstedt tubes (ref 72.693.005). A ∼ 500 µL suspension of cold “acid-washed” beads (Sigma G8772) was added and the tubes were shaken in a PreCellys machine (Bertin Technologies) at 6800 rpm along during two cycles, each cycle comprising 10 seconds ON, 30 seconds OFF and 10 seconds ON. Tubes were placed on ice during 5 min between the two cycles. The lysate was recovered by piercing the bottom of the tube with a heated needle, placing it in a hemolysis tube, and spinning for 1 min at 2500 rpm, 4°C. Total protein amounts in the recovered lysates were quantified using the Bradford-based Bio-Rad Protein Assay (#500-0006) and samples were adjusted to contain 50 µg of proteins in 12 µL. Prior to electrophoresis, 2.5 µL of loading mix 5X (0.5 M Tris pH6.8, 32 % glycerol, 8 % SDS, 8 % DTT, 0.03 % bromophenol blue) was added and tubes were incubated for 5 min at 95°C. Proteins were separated by SDS-PAGE on a 12% acrylamide gel, and transferred in semi-dry conditions to a 0.45 µm nitrocellulose membrane which was then washed in TBS+Tween 0.02% during 10-15 min at room temperature, blocked by incubation in TBS+5% milk for one hour and washed again three times 10 min in TBS + Tween 0.02%. To quantify relative loadings, we incubated the membrane with anti-GAPDH primary antibody (GA1R, Thermofisher MA5-15738) diluted at 1/5000, washed it three times 10 min in TBS + Tween 0.02%, incubated it with HRP-coupled secondary antibody (GE Healthcare NA9310) diluted at 1/5000, washed it three times 10 min in TBS + Tween 0.02%, and revealed it by the ECL chemiluminescent reaction (Cytiva RPN 2232) which was quantified on a Chemidoc reader (BioRad). To quantify LiCre-V5 and Cre-V5 proteins, the membrane was then washed in TBS + Tween 0.02%, incubated with anti-V5 primary antibody (BioRad MCA1360) diluted at 1/1000 for 2 hours at room temperature, washed three times 10 min in TBS + Tween 0.02%, incubated with HRP-coupled secondary antibody (Cytiva NA9310) diluted at 1/5000, washed three times 10 min in TBS + Tween 0.02%, and processed again for ECL and quantification.

For ChIP, aliquoted fixed cells were thawed on ice and resuspended in 300 µL lysis buffer composed of 0.1% w/v sodium deoxycholate, 1 mM EDTA pH8, 50 mM HEPES-KOH pH7.5, 140 mM NaCl, 1% w/v Triton X-100 and proteinase inhibitors (Sigma P2815). Tubes were mixed by pipeting. To disrupt cells, we added 500 µL of cold acid-washed glass beads (Sigma G8772) and placed the tubes on a PreCellys apparatus. Tubes were shaken at 6800 rpm through 2 cycles, each cycle consisting of 10 s ON, 10 s OFF, 10 s ON. Tubes were placed for 1 min on ice between the two cycles. Cell lysates were recovered and sonicated on a Covaris apparatus with the following settings: P 140 W, duty factor 5%, 200 bursts per cycle, 15 min. Debris were eliminated by centrifugation at 14000 rpm 4°C. We then added lysis buffer to reach a volume of 1 mL. After vortexing, 200 µL were used to control sonication efficiency on agarose gels, 750 µL were used for immunoprecipitation (IP) and 50 µL were kept as “inputs”. For IP, anti-V5 monoclonal antibody (BioRad MCA1360) was captured on Dynabeads pA (Life Technologies 10002D) and 35 µL of this preparation were added to the sample, tubes were placed overnight at 4°C on a rotating wheel. In parallel, the “input” samples were also placed overnight at 4°C. The following day, 50 µL of lysis buffer were added to the “input” samples, followed by 35 µL of naked (i.e. without antibody) pA Dynabeads, and this mixture was incubated for 1 hour at 4°C on a rotating wheel. For IP and input samples, beads were washed three times in 0.5 mL of washing buffer I (20 mM TrisHCl pH8, 150 mM NaCl, 2 mM EDTA, 1% Triton X-100, 0.1% SDS), once in 0.5 mL of washing buffer II (20 mM TrisHCl pH8, 500 mM NaCl, 2 mM EDTA, 1% Triton X-100, 0.1% SDS), once in 0.5 mL of washing buffer III (10 mM TrisHCl pH8, 1 mM EDTA, 0.5% deoxycholate, 1% Igepal, 25 mM LiCl) and twice in 0.5 mL of TE8 buffer (10 mM TrisHCl, 1 mM EDTA pH8). For elution, beads were resuspended in 130 µL of Elution Buffer (30 mM TrisHCl pH8, 2 mM EDTA, 1% SDS, 0.35 mg/mL proteinase K), incubated at 65°C for 5 to 6 hours and briefly vortexed every hour. Eluates were transferred to fresh tubes, beads were further incubated in 20 µL of 1 M Tris pH5.5 to generate a second eluate that was also transferred and added to the first. DNA was further purified on NucleoSpin^®^ Gel and PCR Clean-up Columns (Macherey Nagel) and finally eluted in 23 µL ultra-pure water. Real-time qPCR were run on a Rotor-Gene Q thermocycler from QIAGEN using primer pairs (1R69,1R70), (1R71,1R72), and (1R73,1R74) corresponding to probes mentioned in Fig. 1c as P1, P2 and P3, respectively. Unrelated control loci were quantified using primer pairs (1H97,1H98) located in *HIS5*, (1E10,1E11) located in *AHA1* and (1B36,1B37) located in *iSWH1*. We estimated relative quantification (ChIP/INPUT) by computing the NRQ ratio described in [35]. This ratio follows *NRQ = (E^ΔCt^)/R*, where *E* is the amplification efficiency of the target amplicon of interest (for example P1), *ΔCt* is the difference in Ct values for this amplicon between ChIP and INPUT, and *R* is the geometric mean of three reference *E_i_^ΔCti^* values, with *E_i_*being the amplification efficiency of reference *i* and *ΔCti* being the difference in Ct values for reference *i* between ChIP and INPUT. This way, every ratio was normalized by all three references *i* ϵ {*HIS5*, *AHA1, iSWH1}*.

### Analysis by surface plasmon resonance (SPR)

We conducted experiments on a Biacore™ T200 (Cytiva) apparatus equipped with Biacore T200 control software v3.2.1, using streptavidin-coated Series S Sensor Chips SA (Cytiva). We conditioned the chip according to manufacturer’s recommendations (three consecutive 1 min injections of 1 M NaCl in 50 mM NaOH). After conditioning, we used, for all experiments, a running buffer composed of 20 mM Tris pH7.4, 50 mM KCl, 150 mM NaCl, 10 mM MgCl2, 1 mM DTT, 0.05% P20 surfactant (Cytiva). The buffer was filtered on a 0.22 µm MCE membrane (Millipore) before use. Recorded signals were expressed in Response Units (RU) corresponding to a rise in SPR signal resulting from protein/DNA interactions. We prepared ligands by annealing complementary oligonucleotides corresponding to full *loxP* (1Y68/1Y69), or half *loxP* (1Y70/1Y71) or a random sequence (1Y72/1Y73). For each pair, one of the oligonucleotides contained a biotin group at its 5’ end (Supplementary Table S3). Each sensor chip contained 4 flow cells (channels). We first used these channels independently to capture DNA ligands: 32.1 RU of control DNA in channel 1 of the chip, 31.6 RU of half *loxP* DNA in channel 2, 10.6 RU of control DNA in channel 3 and 10.8 RU of *loxP* DNA in channel 4. For kinetic measurements, we then used series of two consecutive channels: FC2-1 (half *loxP)*, and FC4-3 (full *loxP*). Temperature of the chip was set at 25°C. Samples were kept in the Biacore chamber at 10°C. Flow rate was set at 100 µL/min. To measure association, we injected Cre or LiCre for 90 seconds at concentrations varying from 0.3125 to 10 nM prepared as serial two-fold dilutions. We then monitored dissociation for 400 seconds. We applied a regeneration step between each sample to ensure that all detectable Cre or LiCre molecules were dissociated from DNA. These steps consisted of two 30 s injections of 2 M MgCl_2_ in 0.5 M NaCl for full *loxP* kinetics and one 60 s injection of 2 M NaCl for half *loxP* kinetics. Sensorgram artefacts were corrected by subtracting a control injection of running buffer.

### Kinetic model of protein:DNA binding

We based this model on the previous work of [11]. For binding to a half-*loxP* site, we assume that the binding of Cre or LiCre is defined by the association (*k_1_*) and dissociation (*k_-1_*) rates and follows the mass-action model (Fig. 3a):

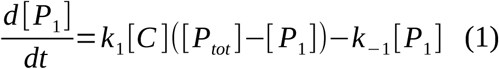

with [*P_tot_*] the total concentration of half-*loxP* sites, [*P*_1_] the concentration of occupied half-*loxP* sites and [*C*] the concentration of Cre/LiCre units. In the SPR protocol, Cre or LiCre proteins are injected at a controlled concentration [*C_i_*]. They diffuse within the biosensor chip and associate with half-*loxP* sites attached to the surface (Fig. 2). This leads to the following equation of evolution for [*C*] :

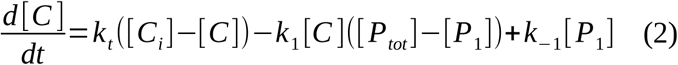

with *k_t_* the coefficient of mass transport within the device.

For binding to a full *loxP* site, we assume a two-step process (Fig. 3a): one unit first binds with the same kinetic rates as for the half-*loxP* site (*k_1_*, *k_-1_*), the second unit then associates with possibly different kinetics (*k_2_*, *k_-2_*). This way, putative cooperative effects can be captured. This leads to the following mass-action model:

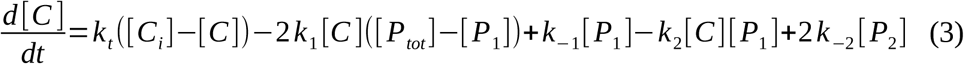

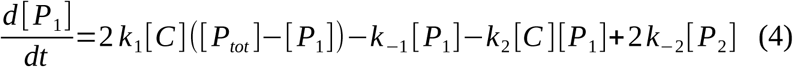

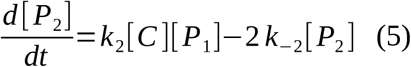

with [*P*_1_] the concentration of half-occupied *loxP* sites and [*P*_2_] the concentration of fully-occupied sites.

The output signal [*B*](*t*) of SPR experiments, given in Response Units (RU), effectively measures the total mass of Cre/LiCre units bound to DNA sites at a given time *t* and is thus proportional to [*P*_1_] in the half-*loxP* experiments, noted ([*B*]*^h^* (*t*)≡*α* [*P*_1_]*^h^*) with superindex *h,* and to [*P*_1_]+2[*P*_2_] in the full-*loxP* experiments, noted ([*B*]*^f^* (*t*)≡*α* ([*P*_1_]*^f^* +2[*P*_2_]*^f^*)) with superindex *f*. The maximal values for [*B*](*t*) are termed *RU_max_* and are defined as 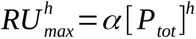 and 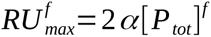. Each experiment consists in two successive steps: an association phase (duration *t_a_*) where the protein is injected ([*C_i_*]≡[*C* _0_]>0 for 0≤*t* ≤*t_a_*) and a dissociation phase in the absence of new injected material ([*C_i_*]=0 for *t*≥*t_a_*). To compare quantitatively model predictions with experiments, for a given experimental condition (Cre or LiCre, half- or full-*loxP* site, [*C* _0_]) and a given set of model parameters (*k_1_*, *k_-1_, k_2_*, *k_-2_, k_t_*, *α, [P_tot_]*), we numerically solved the systems of ordinary differential equations Eqs. (1-2) or Eqs. (3-5) with initial conditions {[*C*],[*P*_1_], [*P*_2_]}={0,0,0} and then computed the corresponding SPR-like output signal (see above).

We fitted model parameters as follows. For each type of molecule (Cre or LiCre) and site (half- or full-*loxP*), experimental data is available for seven values of [*C* _0_] and a fixed *t_a_* value of 90 seconds (Fig. 2). For a given set of parameters (*k_1_*, *k_-1_,k_2_*, *k_-2_, k_t_*, *α, [P_tot_]),* we simulated all experimental conditions (*n*=14) and computed a chi-square-like score defined as:

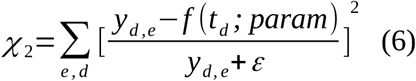

with *y_d,e_* the SPR signal observed at time *t_d_* (∈[0 : 0.1: 340] *sec*) during experiment *e*, *f(t_d_;param)* the corresponding model prediction and ε=10 a meta-parameter that grossly estimates the data uncertainty. Note that one score was defined for the Cre dataset and another for LiCre. The parameters for each type of molecule were fitted independently by minimizing the corresponding *χ* _2_ using a trust region reflective algorithm (function *least_squares* in scipy) starting from initial parameter values randomly sampled within acceptable ranges (Supplementary Table S4). We repeated the minimization 1,000 times per type of molecule and stored, for each fitting round, the optimal set of parameters and the minimum score. The distribution of minimum scores showed that some rounds of minimization only reached a local minimum and did not fully optimize the fit (Figs. 2c-f and 3b-c). We discarded them from the analysis by setting an arbitrary threshold above the first mode of the distribution, keeping only 541 and 859 sets of fitted parameters for Cre and LiCre, respectively.

### Quantification of LiCre-*loxP* recombination in yeast

The yeast assay presented here was performed as previously described [1]. Yeast reporter strains were transformed with the P_met17_-LiCre plasmid pGY466, or plasmid pGY44 (empty vector) or P_met17_-Cre plasmid pGY502 (positive control). For each strain-plasmid combination, three independent transformants were used as biological replicates. For each replicate, a fresh colony was used to inoculate 4 mL of synthetic selective medium lacking methionine (in order to induce LiCre expression) with no particular protection against ambient light. After 18 hours, the saturated culture was transferred to two 96-well polystyrene flat-bottom sterile plates (100 µL per well); one plate was illuminated with the indicated conditions while the other plate was kept in the dark at the same temperature. We used two devices for photoactivation. The first device was a 50 W spot (Neptune-LED, Grenoble, France) equipped with a 450 nm LED connected to a DMX controller, which we placed above a thermostated platform. The second device was a homemade aluminium box which was equipped (by Neptune-LED, Grenoble, France) with a ceiling of LED strips (450 nm) connected to a DMX controller. This box was placed into an incubator at 30°C. Both devices were controlled from a laptop computer via the QLC+ software. After illumination, cells from the two plates were diluted into fresh synthetic medium in deep-well plates and incubated at 30°C for 4 hours to allow expression of the fluorescent protein. We then analyzed cells by flow cytometry either immediately, or the day after. In this latter case, cells were either kept in PBS + 1 mM sodium azide or fixed with 2% PFA before they were analyzed the following day. We then analyzed the raw flow-cytometry data using custom R scripts as previously [1]. No statistical power analysis was done prior to the experiment, no masking of samples IDs was applied during the experiments, and no data randomization was used.

### Kinetic model of recombination

For recombination to happen, we assume that (i) both *loxP* sites should be fully occupied by two LiCre units each and that (ii) at least *x* among 4 LiCre units should be currently in the activated state. If we neglect the backward, re-insertion reaction (see main text), the probability *p* for a cell to have not recombined yet follows the evolution equation:

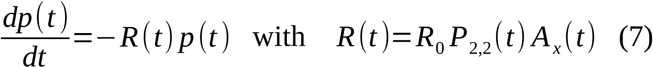

where *P*_2,2_ (*t*) is the probability at a given time *t* that condition (i) is effective, *A_x_* (*t*) is the probability that condition (ii) is realized and *R*_0_ is the recombination rate when both conditions (i) and (ii) are fulfilled. Note that *R*_0_ encompasses the rate of DNA looping between the two *loxP* sites, the rate of synapse formation and the actual rate of recombination when the synapse is formed and functional. From Eq. 7, the fraction *ϕ(t)* of cells having recombined at time *t* is given by

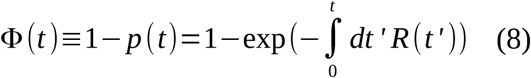

where we assumed that initially (*t=0*) no cells have recombined (*p(0)=1*).

#### Occupation of loxP sites

We assume that binding of LiCre at one *loxP* is independent from binding events at the other *loxP* site, which we write as:

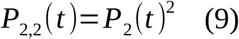

with *P*_2_(*t*) the probability that one site is fully occupied, which is given by Eqs. (4-5) by replacing [*P*_1_] and [*P*_2_] by *P*_1_ (*t*) and *P*_2_ (*t*) (the probabilities that the site is occupied by one and two LiCre units, respectively), and by setting [*C*] as a constant equal to the average nuclear concentration of LiCre.

Since light does not impact LiCre binding (Fig. 1) and binding/unbinding of LiCre is much faster (∼ seconds, Fig. 3f) than the recombination step (minutes), we assume that occupation of *loxP* sites is at steady-state. Finding the stationary solutions of Eq. (4-5) and using Eq. (9) leads to

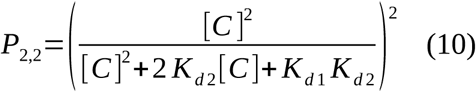

#### Photo-activation of individual LiCre units

Assuming that the activation of one LiCre unit is independent from the state of the 3 other units bound to the two *loxP* sites,

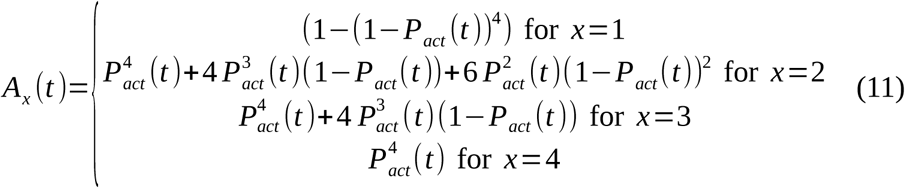

with *P_act_* (*t*) being the probability that one LiCre unit is activated. The kinetics of activation/inactivation is supposed to follow a simple two-state model:

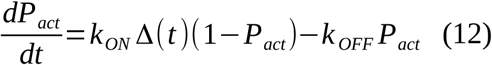

where *k_ON_* is the activation rate when light is ON and depends on the lighting intensity (see below), Δ*(t)* is a time-dependent binary variable that is equal to 1 when the light is ON and 0 otherwise, *k_OFF_* is the deactivation rate. For a periodic illumination defined by a period *T* and a duty cycle *dc*, that represents the fraction of *T* during which the light is ON, Eq.(12) can be solved analytically:

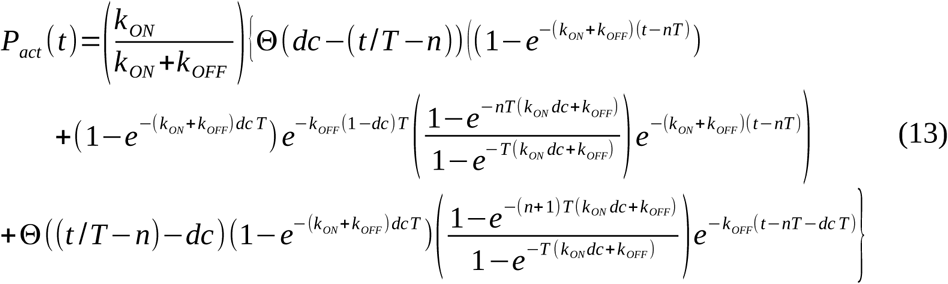

with *n*=⌊*t* /*T* ⌋ and Θ(*u*) being the Heaviside function.

#### Choice of parameters, integration and fitting

For a given set of parameters *(R_0_, K_d1_, K_d2_, [C], k_ON_, k_OFF_, T, dc),* the time evolution of *ϕ(t)* can be computed using Eqs. (7,8,10,11,13). To compare it to experimental data, we fix parameters for which a value was known. We chose the *K_d1_, K_d2_* values that were inferred from SPR experiments on LiCre (Fig. 3f). For *k_ON_*, we used the estimate of [13] who computed the activation rate of a LOV2 domain as *k_ON_*=0.26 *S F* with *S=4.3 10^−17^ cm^2^* the surface of LOV2, *F=(E λ 10^13^)/2 cm^−2^s^−1^* the flux of incoming photons, *λ* the light wavelength (in *nm*), and *E* the power of the light in *mW.cm^−2^.* In our case, this led to *k*_ON_ ≈(*E* /35) *s*^−1^. The concentration of LiCre in the yeast nucleus is not known. For simplicity, we fixed it at *[C]=1 nM* which represents about 100 molecules per nucleus. *E*, *T* and *dc* are defined by the illumination conditions of each experiment.

To fit the remaining parameters (*x, R_0_* and *k_OFF_*), we considered 4 sets of experiments listed in Supplementary Table S5, each set corresponding to a coherent series (proximal dates, same operator and comparable conditions): experiment 8 (Fig. 5d), experiments 1 and 2 (Fig. 5e), experiments 3 to 7 (Fig. 5f) and experiment 9 (Fig. 5g). For each set of experiments, we systematically varied *x* from 1 to 4, *R_0_* between 5.10^−5^ and 1.5 10^−3^ *s^−1^* and *k_OFF_* between 8.10^−4^ and 2.5 10^−2^ *s^−1^*, we computed the corresponding model prediction for *ϕ(t)* and we estimated a chi-square-like score (similar to Eq. 6) where experimentally-measured recombination efficiencies were normalized by a factor 0.87 to account for the frequency of LiCre-plasmid loss (see main text). For each set *e* of experiments and each *x* and *k_OFF_* values, we estimated the minimum chi-square-like score 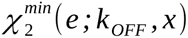 and the corresponding *R_0_(e)* value. We then estimated the value of *x* and *k_OFF_* that minimized 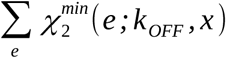. For each *x* value, optimal *{R_0_(e;x)}* and *k_OFF_(x)* values are shown in Table 1.

Note that we fitted the model by imposing the same values of *x* and *k_OFF_* for every experiments. In contrast, *R_0_* was estimated for each experiment in order to capture the observed batch-to-batch experimental variability (which was presumably due to changing environmental conditions such as temperature, see main text). Note also that our arbitrary choice for *[C]* does not impact the quality of the fit nor the best values for *x* and *k_OFF_*, because changing its value only changes *P*_2,2_ (Eq.10) which acts as a normalizing factor in Eq.(7). Nonetheless, the choice of *[C]* may impact on the fitted values of *R_0_(e;x)* as follows: 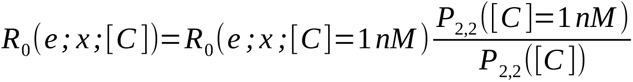 which has a rather slow dependency in *[C]* and thus the fitted value would be rather similar unless *[C]<<1 nM* which is unlikely (see Discussion below).

To separately estimate *k_OFF_* and *R_0_*in WT LiCre and in the T418S mutant, we applied a similar strategy as described above but using only dataset 13 (Supplementary Table S5). For each variant (WT or T418S) and each *x* value, we computed a chi-square score over a 2D-grid of parameters for *k_OFF_*and *R_0_*, accounting for all the experimental data of the variant. Parameters that minimize this score are given in Supplementary Table S7.

### Relation between *R_0_* and kinetic parameters previously-estimated for Cre

In [23], authors fitted *in-vitro* FRET measurements by a three-state model of intra-molecular Cre-*loxP* recombination (Fig. 6A of their publication). They estimated the rate of synapse formation from the unlooped state *k_3_*, the rate of synapse unfolding *k_-3_* and the rate of recombination catalysis *k_-4_*. They found that *k_-4_* was the rate-limiting step in the two cases investigated (inter-*loxP* distances of 3044 and 870 bp). In our model, *R_0_* encompasses all these different reactions. In the limit *k_-4_<< k_3_, k_-3_*, we can approximate *R_0_* by the probability for a non-edited DNA molecule to be in the synaptic state, multiplied by the catalysis rate, giving *R*_0_=*k*_−4_ (*k*_3_)/(*k*_3_ +*k*_−3_). We used this formula to compare our *R_0_* values with corresponding estimates from Table 2 of [23] (see Discussion).

### Intracellular LiCre concentration

The effective concentration of ‘free’ recombinase available to conduct DNA recombination in live cells is unknown. It is, however, related to the average intracellular concentration of LiCre, which we anticipate to be largely above 1 nM for the following reasons. In our conditions, LiCre expression was driven by the P_MET17_ promoter and its level is therefore likely comparable to the level of Met17p. In a previous study based on LC-MS, this protein was estimated at ∼120,000 copies per yeast cell (equivalent to 4 μM if we assume a cell volume of 50 µm^3^), an abundance 1.6 times higher than the average abundance of the proteins quantified in this study (Data S1B of [36]). Another study based on fluorescently-tagged proteins [37] estimated an average concentration of 12,100 copies per cell (0.4 µM) per protein. So, unless LiCre is particularly unstable in yeast, its concentration should be 2 or 3 orders of magnitude above 1 nM.

## ACKNOWLEDGEMENTS

This work has benefited from the expertise of the Protein Science (Virginie Gueguen-Chaignon and Aurélie Thibaut) and Flow Cytometry facilities of the SFR Biosciences Lyon. We thank Fabien Duveau, Mirko Francesconi and Aurèle Piazza for fruitful discussions, Fabien Duveau, Mirko Francesconi and Valérie Robert for critical reading of the manuscript, five anonymous reviewers for their comments, Pascal Bernard for a plasmid carrying V5-epitope repeats, Jean-Philippe Robin for advice on recombinant proteins and developers of R, Bioconductor, QLC+, numpy, git and Ubuntu for their software.

## FUNDING

This work was supported by CNRS under the “MITI 80 Prime” program grant READGEN, the Fondation ARC pour la recherche contre le cancer, ANR Grant Opto4D ANR-23-CE12-0039, a funding for interdisciplinary innovations from the Institut RhôneAlpin des Systèmes Complexes (IXXI) and the Federation for Systems Biology in Lyon (BioSyL) and a funding for technological developments from the SFR Biosciences Lyon.

## DATA AVAILABILITY

Supplementary Data contain the source data of all figures (Supporting Dataset), computational codes implementing the models of LiCre:DNA binding and recombination reaction, which are also made available at https://github.com/physical-biology-of-chromatin/licre-optogenetic-kinetics/, as well as Supplementary Tables. Plasmids used for the production of recombinant LiCre (pET-LiCre = pGY611) or Cre (pET-Cre = pGY709) are made available from Addgene (https://addgene.org) under accession ID 219797 and 219798, respectively.

## AUTHORS CONTRIBUTION

Designed and supervised the study: D. J. for modeling, G. Y. for experiments (except SPR).

Designed and performed the SPR experiment: A. Duf, C.M. and G. T.

Constructed plasmids and strains: A. Dum, H. D-B., G. T.

Generated datasets 1-8 and 10-11: H. D-B.

Generated dataset 9: C. D-K.

Performed ChIP and WB experiments: H. D-B.

Constructed the DNA-binding model: T. B-L., E. C. and D. J.

Fitted parameters of the DNA-binding model: D. J. and G. Y.

Constructed and analyzed the recombination model: T. B-L., E. C. and D. J.

Fitted parameters of the recombination model: D. J. and L.T.

Analyzed flow-cytometry data: G. Y.

Designed and constructed LED box chassis: F. V.

Interpreted results: A. Duf., D. J., C. M., G. T., G. Y.

Made the figures: G. Y.

Wrote the paper: D. J., G. Y. with corrections from C. M. and inputs from all the authors.

Read and approved the manuscript: all authors

## Notes

### Competing Interest Statement

The authors have declared no competing interest.

### Summary of Updates

- Several control experiments. - Experimental analysis of LOV2 mutants affecting its light cycle (Fig 6). - Transfer of supp figs into main figs.

