## Supplementary Tables for "Kinetic properties of optogenetic site-specific DNA recombination by LiCre-*loxP*"

**Supplementary Table S1.** List of plasmids used in this study.

| ID | Name | System | Type | Description | Reference |
|---|---|---|---|---|---|
| pGY44 | pRS314 | yeast | Centromeric | Empty vectorBrachmann et al. 1998 | Brachmann et al. 1998 |
| pGY466 | Pmet17-LiCre | yeast | Centromeric | LiCre expression under Pmet17 promoter | Duplus-Bottin et al. 2021 |
| pGY472 | pGY472 | yeast | Integrative | HO-L:Ptef-loxP-KlLEU2-Tadh1-loxP-mCherry-Tcyc1:HO-R | Duplus-Bottin et al. 2021 |
| pGY502 | Pmet17-Cre | yeast | Centromeric | Cre expression under Pmet17 promoter | Duplus-Bottin et al. 2021 |
| pGY537 | pGY537 | yeast | Integrative | pISLys2:Ptef-loxP-KlLEU2-Tadh1-loxP-GFP-URA3 | Duplus-Bottin et al. 2021 |
| pGY605 | Pmet17-LiCre-V5 | yeast | Centromeric | LiCre-V5 expression under Pmet17 promoter | This work |
| pGY607 | Pmet17-Cre-V5 | yeast | Centromeric | Cre-V5 expression under Pmet17 promoter | This work |
| pGY611 | pET-LiCre | E. coli | IPTG expression | For production of recombinant LiCre in *E. coli* | This work |
| pGY618 | pGY618 | yeast | Integrative | pISLys2:Ptef-loxP-KlLEU2-Tadh1-loxP-mCherry-URA3 | This work |
| pGY621 | pGY621 | yeast | Integrative | pISLys2:Tadh1-loxP-mCherry-Tcyc1-URA3 | This work |
| pGY709 | pET-Cre | E. coli | IPTG expression | For production of recombinant Cre in *E. coli* | This work |
| pGY775 | pLiCre-Q513L | yeast | Centromeric | LiCre-Q513L expression under Pmet17 promoter | This work |
| pGY776 | pLiCre-V416L | yeast | Centromeric | LiCre-V416L expression under Pmet17 promoter | This work |
| pGY777 | pLiCre-F494I,L496S | yeast | Centromeric | LiCre-F494I,L496S expression under Pmet17 promoter | This work |
| pGY778 | pLiCre-V416T | yeast | Centromeric | LiCre-V416T expression under Pmet17 promoter | This work |
| pGY779 | pLiCre-N425Q,I427V | yeast | Centromeric | LiCre-N425Q,I427V expression under Pmet17 promoter | This work |
| pGY780 | pLiCre-T418S | yeast | Centromeric | LiCre-T418S expression under Pmet17 promoter | This work |
| pGY781 | pMet-LiCre-FMN1 | yeast | Centromeric | LiCre expression under Pmet17 promoter and FMN1 expression under Padh1 promoter | This work |

**Supplementary Table S2.** List of yeast strains used in this study.

| ID | Background | Genotype[(i)] | Source |
|---|---|---|---|
| OAy470 | W303 | *MATa ade2-1 his3-11,15 leu2-3,112 trp1-1 ura3-1 bar1::hisG* | Aparicio et al. 1997 |
| GY855 | S288c (BY) | *MATa leu2Δ0 trp1Δ63 ura3Δ0* | Duplus-Bottin et al. 2021 |
| GY1761 | S288c (BY) | *MATb his3Δ200 leu2Δ1 trp1Δ63 ura3Δ0 hoΔ::loxKlLEU2loxGFP* | Duplus-Bottin et al. 2021 |
| GY2214 | S288c (BY) | *MATa/MATb ADE2/ade2Δ::hisG his3Δ200/HIS3 leu2Δ0/leu2Δ1 LYS2/lys2Δ::loxKlLEU2loxGFP trp1Δ63/trp1Δ63 ura3Δ0/ura3Δ0 HO/hoΔ::loxKlLEU2loxmCherry* | Duplus-Bottin et al. 2021 |
| GY2416 | S288c (BY) | *MATa leu2Δ0 lys2Δ::Tadh1-loxP-mCherry trp1Δ63 ura3Δ0* | This work |
| GY2450 | S288c (BY) | *MATa leu2Δ0 lys2Δ::loxKlLEU2lox-mCherry trp1Δ63 ura3Δ0* | This work |
| GY2517 | S288c (BY) | *MATa/MATb his3D200/HIS3 leu2D0/leu2D0 LYS2/lys2Δ::loxKlLEU2loxmCherry trp1Δ63/trp1Δ63 ura3Δ0/ura3Δ0 HO/hoΔ::loxKlLEU2loxGFP* | This work |
| GY2752 | W303 | *MATa ADE2 his3-11,15 leu2-3,112 trp1-1 ura3-1 bar1::hisG* | This work |
| GY2753 | W303 | *MATa ADE2 his3-11,15 leu2-3,112 trp1-1 ura3-1 hoΔ::loxKlLEU2LoxmCherry bar1::hisG* | This work |

(i): *MATb* corresponds to *MAT***α** *(alpha)*

**Supplementary Table S3.** List of oligonucleotides used in this study.

| ID | Sequence (5' to 3') |
|---|---|
| 1B36 | TGCTGAGTTTTTGCGCATCAAT |
| 1B37 | TTTTGGTGCACGTTCGCT |
| 1E10 | CGTGAATAACCCAAATAACTGG |
| 1E11 | GCATACTTTTTATCCTTCACCG |
| 1H97 | GGGGTGACAATGTCTTGGCAAA |
| 1H98 | ACATCCGGAACAGCCTAATTCG |
| 1R69 | TTATCGATGCATGCCTGCAGGT |
| 1R70 | TTGACAGCCTTGGCGATAGCAT |
| 1R71 | TTGACAGCCTTGGCGATAGCAT |
| 1R72 | CGCCTTTGGTCACCTTCAATTTGG |
| 1R73 | TTTGAAGACGGTGGGGTTGTAACG |
| 1R74 | GCATGACAGGACCATCAGAAGGAA |
| 1Y68 | biotin-CGGAATTCACAACTTCGTATAATGTATGCTATACGAAGTTATGGATCCGC |
| 1Y69 | GCGGATCCATAACTTCGTATAGCATACATTATACGAAGTTGTGAATTCCG |
| 1Y70 | biotin-CGGAATTCAGATCTATAACTTCGTATAATGTGTCTAGAGGATCCGC |
| 1Y71 | GCGGATCCTCTAGACACATTATACGAAGTTATAGATCTGAATTCCG |
| 1Y72 | biotin-CGGAATTCAGATCTACATATGTGATATCTAAGCTTATCTAGAGGATCCGC |
| 1Y73 | GCGGATCCTCTAGATAAGCTTAGATATCACATATGTAGATCTGAATTCCG |
| 1B36 | TGCTGAGTTTTTGCGCATCAAT |
| 1B37 | TTTTGGTGCACGTTCGCT |

**Supplementary Table S4.** Boundaries of initial parameter values that were used to fit the DNA-binding model to experimental SPR data.

| | **Cre** | | **LiCre** | |
|---|---|---|---|---|
| **Parameter** | **Lower bound** | **Upper bound** | **Lower bound** | **Upper bound** |
| $\alpha$ | 1 | 2.5 | 1 | 2.5 |
| $RUmax_h$ | 36 | 44 | 54 | 62 |
| $RUmax_f$ | 24 | 30 | 36 | 44 |
| $k_t$ | 0.08 | 1.1 | 0.08 | 1.1 |
| $k_1$ | -3 | 2 | -3 | 2 |
| $k_{-1}$ | -3 | 2 | -3 | 2 |
| $k_2$ | -4 | 2 | -4 | 2 |
| $k_{-2}$ | -4 | 0 | -4 | 0 |

**Supplementary Table S5.** Experimental data sets quantifying LiCre efficiency in yeast cells.

| ID | Date | Strain | DMX Device | Intensity (mW/ $cm^2$) | Duration (min) | Regime | Temp. (°C) | Source | Related figures |
|---|---|---|---|---|---|---|---|---|---|
| 1 | 2020-11 | GY2214 | Spot | 35 | 60 | Period: 10 s or 2 min<br>Duty: 0, 1, 5, 10, 50 or 100% | Room Temp. | Duplus-Bottin 2021 | 5e |
| 2 | 2020-09&10 | GY2214 | Spot | 1.5 | 60 | Period: 2 min<br>Duty: 0, 5, 10, 50, 75 or 100% | Room Temp. | Duplus-Bottin 2021 | 5e |
| 3 | 2021-10 | GY2517 | Box | 35 | 7,20,60, or 180 | Period: 10 s<br>Duty: 10% | 30 | This work | 4g, 5f |
| 4 | 2022-03-14 | GY2517 | Box | 35 | 20 or 60 | Period: 20 s<br>Duty: 10% | 30 | This work | 4g, 5f |
| 5 | 2022-03-21 | GY2517 | Box | 35 | 7, 20 or 60 | Period: 5 s<br>Duty: 10% | 30 | This work | 4g, 5f |
| 6 | 2022-04-04 | GY2517 | Box | 35 | 7, 20 or 60 | Period: 60 s<br>Duty: 10% | 30 | This work | 4g, 5f |
| 7 | 2022-04-12 | GY2517 | Box | 35 | 7, 20 or 60 | Period: 60 s<br>Duty: 33.3% | 30 | This work | 5f |
| 8 | 2019-07-11&12 | GY2214 | Spot | 35 | 10, 20, 30, 60, 90, 180, 240 | Period: 10 s<br>Duty: 5% | Room Temp. | This work | 4e, 5d |
| 9 | 2023-05-03 | GY2753 | Spot | various | 60 | Period: 7 s<br>Duty: 28.6% | 30 | This work | 4h (red), 5g |
| 10 | 2020-07-03 | GY2214 | Spot | 35 | 60 | Period: 10 s<br>Duty: 5% | 20, 25, 30, 37 | This work | 4f |
| 11 | 2024-04-12 | GY2753 | Spot | various | 60 | Period: 7 s<br>Duty: 28.6% | 30 | This work | 4h (black) |
| 12 | 2024-10-18 | GY2517 | Box | 35 | 60 | Period: 10 s<br>Duty: 10% | 30 | This work | 4c, 4d, 6a |
| 13 | 2025-01-17 | GY2517 | Box | 35 | 7,20,60 | Period: 10 s or 60 s<br>Duty: 0, 10, 20 or 50% | 30 | This work | 6b |

**Supplementary Table S6.** Assay evaluating frequency of plasmid loss

| Culture | Dilution | CFUs on selective SD-W plates | CFUs on non-selective SD plates | Total CFUs on selective | Total CFUs on non-selective | Loss in this culture (%) | Loss (%) |
|---|---|---|---|---|---|---|---|
| A | A1 | 192 | 216 | 972 | 1083 | 10.25 | 21.45 |
| | | 198 | 226 | | | | |
| | | 237 | 233 | | | | |
| | A2 | 115 | 148 | | | | |
| | | 114 | 125 | | | | |
| | | 116 | 135 | | | | |
| B | B1 | 226 | 293 | 859 | 1190 | 27.82 | |
| | | 174 | 218 | | | | |
| | | 182 | 271 | | | | |
| | B2 | 92 | 145 | | | | |
| | | 91 | 144 | | | | |
| | | 94 | 119 | | | | |
| C | C1 | 198 | 271 | 889 | 1206 | 26.29 | |
| | | 180 | 234 | | | | |
| | | 200 | 298 | | | | |
| | C2 | 101 | 150 | | | | |
| | | 112 | 122 | | | | |
| | | 98 | 131 | | | | |

**Supplementary Table S7:** Best-fit model parameters using dataset Nb 13 only. The model was fitted on LiCreWT data and on LiCreT418S data separately.

| Parameter | Data set specificity | $x$ = 1 | $x$ = 2 | $x$ = 3 | $x$ = 4 |
|---|---|---|---|---|---|
| $k_{OFF}$ ($10^{-3}$ $s^{-1}$) | LiCre-WT | 79.4 | 39.7 | 19.9 | 8.9 |
| $k_{OFF}$ ($10^{-3}$ $s^{-1}$) | LiCre-T418S | 100 | 62.9 | 39.7 | 17.7 |
| $R_0$ ($10^{-4}$ $s^{-1}$) | LiCre-WT | 2.2 | 2.2 | 2.2 | 2.5 |
| $R_0$ ($10^{-4}$ $s^{-1}$) | LiCre-T418S | 2.2 | 2.5 | 2.8 | 3.2 |
| Fit score[a] | LiCre-WT | 0.097 | 0.097 | 0.099 | 0.102 |
| Fit score[a] | LiCre-T418S | 0.097 | 0.094 | 0.091 | 0.088 |

(a) Chi-square-like score quantifying distance to observations. The lower this score, the better the fit (see Methods).
